# Anatomical White Matter Tracts Span the Cortical Hierarchy to Support Cognitive Diversity

**DOI:** 10.64898/2025.12.20.695692

**Authors:** Joëlle Bagautdinova, Golia Shafiei, Audrey C. Luo, Margaret K. Pecsok, Taylor Salo, Aaron F. Alexander-Bloch, Dani S. Bassett, Margaret E. Gardner, Raquel E. Gur, Ruben C. Gur, Allyson P. Mackey, Bratislav Misic, Tyler M. Moore, David R. Roalf, Russell T. Shinohara, Valerie J. Sydnor, Tien T. Tong, Fang-Cheng Yeh, Russell A. Poldrack, Matthew Cieslak, Theodore D. Satterthwaite

## Abstract

Long-range white matter (WM) tracts support cognition by enabling communication between distant cortical regions, which are organized along a hierarchy defined by the sensorimotor-to-association (S-A) axis. However, it remains unknown how WM tracts are positioned within the cortical hierarchy to support cognition. Here, we show that WM tracts are differentially positioned in the cortical hierarchy to support specific cognitive functions, and that tracts spanning the hierarchy connect regions with greater cognitive diversity. Moreover, tracts situated within the same hierarchical level connect biologically similar regions, while those crossing the hierarchy bridge distinct biological milieux to support diverse cognitive functions. The placement of tracts in the cortical hierarchy also reflects developmental variation in tract microstructure and individual differences in cognition. Together, these findings provide a framework that moves beyond conventional categories of association or projection tracts and links WM tract anatomy to cortical organization, cognitive function, cortical neurobiology, and neurodevelopment. We anticipate that this cortex-anchored framework for describing WM tracts may aid the interpretation of individual differences in WM structure related to development and behavior.

## Introduction

Long-range white matter (WM) tracts were first delineated by post-mortem dissection, and are thought to enable communication among distant brain regions that is essential for complex cognition^1,2^. Higher-order functions rely on spatially distributed regions embedded in a cortical hierarchy defined by the sensorimotor-to-association (S-A) axis^3–7^. This major axis of brain organization also describes variation in diverse biological properties^8^, including intracortical myelination^9^, gene expression^10,11^, laminar differentiation^12–14^, and neurotransmitter receptor density^17^—all of which are thought to be related to WM connectivity^15–23^. However, only sparse work has sought to link canonical WM tracts to cognition or the biological milieu of their cortical endpoints. Here, we sought to delineate how anatomically defined, canonical WM tracts are situated within the broader context of cortical organization to support unique cognitive functions.

This study addresses three major knowledge gaps. First, despite extensive research, methodological limitations have historically made it difficult to study the function of WM tracts based on their cortical connectivity profiles in humans. To date, the relationship between WM connectivity and function has been examined primarily using inter-regional connectomes defined using whole-brain tractography^24,25^. However, inter-regional connectomes do not typically describe which WM tracts enable cortico-cortical connectivity. As such, edges in these connectomes are effectively agnostic to anatomical WM tracts. Additionally, whole-brain tractography approaches that produce structural connectomes suffer from high rates of false positives; large-scale studies across several research groups have suggested that tractography can generate streamlines that do not reflect ground-truth anatomic connections^26^. Together, these limitations challenge the biological plausibility of inter-regional structural connectomes, highlighting the importance of linking the known structure of canonical WM tracts to function.

A second key gap is that there have been only sparse efforts to map canonical WM tracts directly to cognitive functions. Prior work includes conceptual reviews of evidence from post-mortem dissection, histology, and theoretical knowledge of tracts^27^; lesion-based mappings of cognitive deficits^28^; and efforts to project functional activations from task-based fMRI onto WM^29,30^. While informative, these approaches remain limited: conceptual reviews are interpretative rather than empirical; lesion-based studies rely on naturally occurring lesions; and most studies have not used anatomically-validated tracts. Together, these limitations underscore the need to systematically map the cognitive functions associated with anatomical WM tracts.

Third, we lack a clear understanding of how WM tracts align with large scale patterns of cortical organization. Convergent lines of evidence emphasize that the cortex is organized along a dominant S-A axis, spanning from sensorimotor to association regions^3–7^. Since cognitive functions are embedded in this cortical hierarchy^3,5^, understanding how major WM tracts are situated in the hierarchy to support specific cognitive functions is critical. Furthermore, the cognitive profiles of tracts are likely shaped by the neurobiological properties of their cortical endpoints. However, existing evidence is divided on how those WM tract endpoints are arranged: some studies suggest that long-range WM connections preferentially link regions with similar biological profiles^20,21,31–33^, while others report that they instead connect regions with diverse biological properties^34^. This discrepancy reinforces the need to explicitly link WM tracts to the cortical hierarchy and biological properties of their cortical endpoints as a crucial step towards understanding how WM supports brain function^27^.

In the context of these limitations, recent methodological advances have enabled the mapping of WM tract endpoints onto the cortical surface in individual brains, yielding person-specific maps that link tracts to cortex^35^. By aggregating across individuals, these mappings form a population-level tract-to-region connectome that captures both the anatomical WM tracts involved and their cortical endpoints. Importantly, this tract-to-region mapping represents an anatomically-grounded alternative to conventional inter-regional structural connectomes, and provides the foundation for annotating tracts with the cognitive functions and biological properties of their endpoint regions.

Here, we aimed to delineate how anatomical WM tracts are situated within the cortical hierarchy to support cognition. As described below, we integrated several multimodal features to elucidate the spatial, hierarchical, functional, and biological characteristics of the cortical regions that each WM tract connects. We hypothesized that WM tracts would markedly differ in how they connect regions in the cortical hierarchy, conferring varying levels of cognitive diversity or specialization. We also expected that the diversity of tract functions is underpinned by the neurobiological characteristics of their cortical endpoints, where tracts supporting diverse cognitive functions act as bridges across distinct neurobiological milieux. Finally, we evaluated whether the position of a tract in the cortical hierarchy captures meaningful variation in brain development and individual differences in cognition using large-scale data from the Philadelphia Neurodevelopmental Cohort^37^. We hypothesized that WM tract which span the cortical hierarchy would show stronger associations with development and executive function. As described below, this framework provides a unified approach linking the architecture of canonical WM tracts to cortical organization, cognitive function, cortical neurobiology, and neurodevelopment – revealing how tracts differentially span the cortical hierarchy to support unique forms of human cognition. In doing so, our results serve as a practical reference for researchers and clinicians interested in the typical functions associated with specific long-range WM tracts.

## Results

### Functional and neurobiological annotations of WM tracts

To delineate how anatomically defined white matter tracts are situated within the cortex to support cognition, we leveraged three population-level brain data resources. First, we used tract-to-region mappings from population-level tractography in young adults from the Human Connectome Project (HCP-YA, *N*=1,065)^35^, which established links from tracts to cortex and provided the foundation for annotating tracts with cortical properties (**Figure 1a**). This tract-to-region connectome describes the connections of 52 tracts (26 per hemisphere) to 360 cortical brain regions of the HCP-MMP parcellation^9^, resulting in a 360-by-52 matrix (see **Supplementary Table 1** for a list of all tracts, their abbreviations, and classification into projection or association categories). Importantly, this tract-to-region connectome captures both the specific WM tracts along with their endpoints, providing a biologically-grounded alternative to conventional region-to-region structural connectomes that lack tract identity information and are susceptible to high rates of false positives^26^.

**Figure 1.**
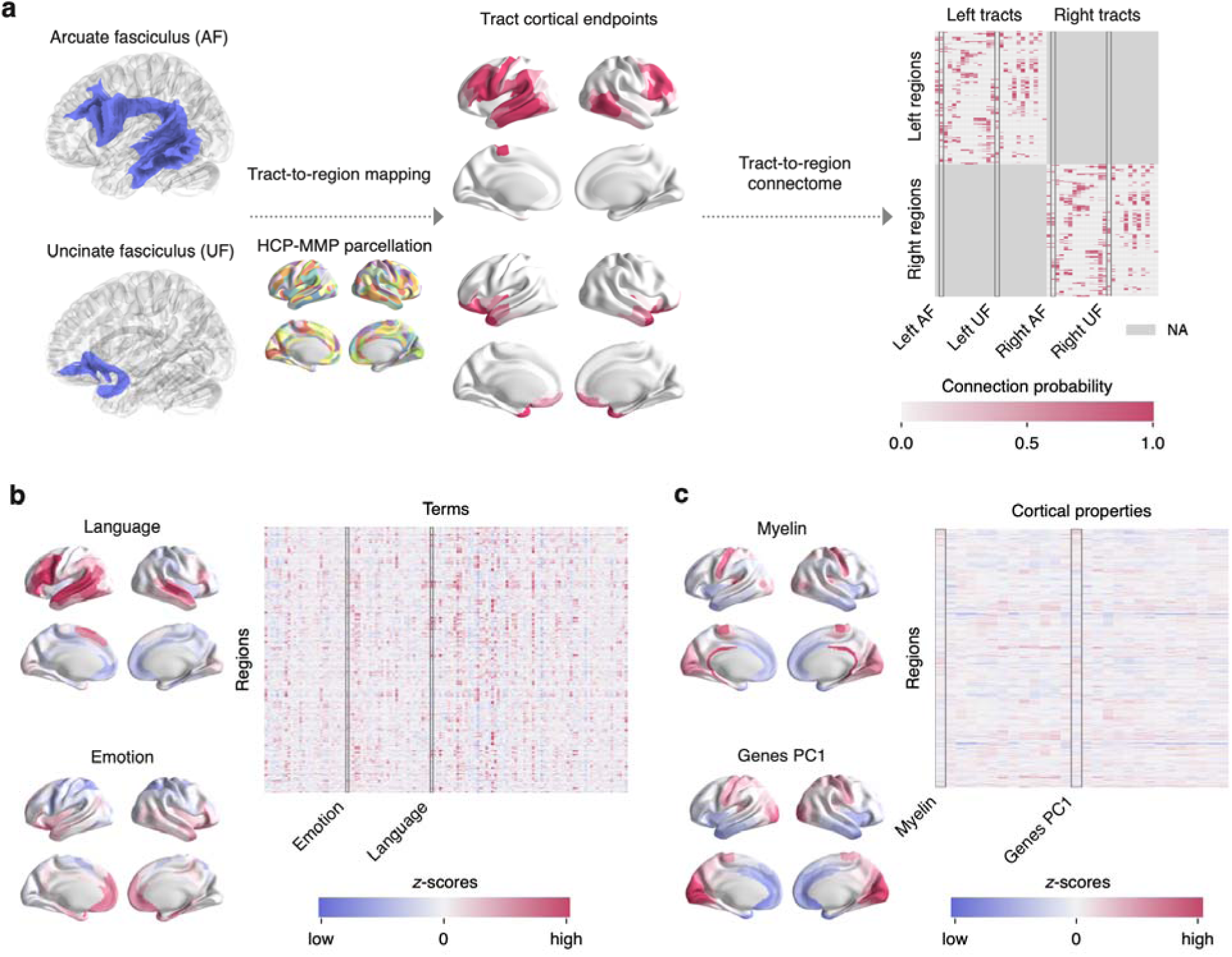
Overview of data types. **a)** The tract-to-region connectome used in this study was generated and described in detail by Yeh, 2022. Briefly, tractography was run in *N*=1,065 HCP-YA participants to reconstruct 52 major WM tracts. For each tract, cortical endpoints were intersected with 360 brain regions from the HCP-MMP parcellation and then aggregated across all participants to create a population-level map representing the probability of connection of the tract to each cortical region. This process is performed for all tracts to obtain a population-level matrix of 360 regions. Because this analysis was focused on within-hemisphere tracts, the matrix elements corresponding to contralateral regions were set to NA. **b)** The *Cognitive Atlas* was used to select 125 meta-analytic cognitive terms within the *Neurosynth* database corresponding to well-defined lower- to higher-order cognitive processes. Left: example activation maps for “Language” and “Emotion”. Right: the resulting cognitive terms matrix of 360 brain regions by 125 cognitive terms, where values correspond to association test *z*-scores quantifying to what extent a given cortical region was more consistently activated in studies that mentioned the term, as opposed to studies that did not mention it. **c)** Biological cortical properties available through *Neuromaps* and *BigBrain*. Left: example maps for intracortical myelination (“Myelin”) and the first principal component of gene expression (“Genes PC1”) across the cortex. Right: the resulting matrix of 360 brain regions by 28 biological cortical properties, where values correspond to *z*-scores of cortical features.

Second, we leveraged prior task-based fMRI results, collected and meta-analyzed via *Neurosynth*^36^, to gain insight into the cognitive profiles of tracts. We drew on the *Cognitive Atlas*^38^ to select 125 cognitive terms within the *Neurosynth* database that corresponded to well-characterized cognitive functions ranging from lower- to higher-order cognitive processes. This resulted in a cognitive terms matrix of 360 brain regions by 125 cognitive terms (**Figure 1b**). The elements of the cognitive terms matrix reflected association test *z*-scores quantifying how consistently a cortical region was activated in fMRI studies that mentioned the term.

Third, to delineate the biological properties of regions connected by specific WM tracts, we selected 28 feature maps from *Neuromaps*^8^ and *BigBrain*^12–14^ based on their relevance to WM anatomy, including intracortical myelination^9^, gene expression^10,11^, laminar differentiation^12–14^, and neurotransmitter receptor density^15,39–57^. This resulted in a matrix of 360 brain regions by 28 biological cortical properties, where elements corresponded to the *z*-scores of cortical features (**Figure 1c**).

In summary, we used the tract-to-region mappings as a bridge between canonical WM tracts and rich information regarding the function and biology of the cortical regions they connect. As reported below, we begin by describing tracts’ spatial embedding in the cortical hierarchy. Next, we link tracts to cognitive functions and quantify tracts’ cognitive diversity. We then situate the cognitive diversity of WM tracts in the context of the biological features of the cortical regions with which they connect. Finally, we move beyond population-average features to investigate how the position of tracts within the cortical hierarchy is related to brain development in youth and individual differences in cognition.

### Tracts vary in their embedding in the cortical hierarchy

While canonical WM tracts enable long-range communication in the brain, their positioning within the brain’s cortical hierarchy remains sparsely described. We hypothesized that the spatial distance between regions connected by a tract would vary as a function of the regions’ position in the cortical hierarchy. Specifically, we expected that short-range tracts would connect cortical regions that were at a similar level of hierarchy, whereas long-range tracts would connect regions at different levels of the cortical hierarchy. As a first step, we sought to characterize the spatial and hierarchical embedding of long-range tracts. Two different inputs were required to do this. First, to assess spatial distance, we used the centroid coordinates of brain regions to generate a region-to-region matrix containing the Euclidean distance between region pairs, reflecting the straight line connecting two regions in 3D space (**Figure 2a**). Second, we quantified cortical hierarchy using the S-A axis^3^ (**Figure 2b**). To quantify the spatial distance spanned by regions connected to each tract, we calculated the mean Euclidean distance between connected region pairs for each tract (**Figure 2c**). To determine the hierarchical positioning of tracts, we evaluated the degree to which each tract spanned the cortical hierarchy by computing the tract’s S-A range (e.g., the difference between the maximum and minimum S-A rank of regions connected to each tract; **Figure 2c**). The association between tract annotations was quantified using Spearman’s rank correlation (*r_s_)*; the significance of the association was tested using a non-parametric permutation test (*p*_perm_). We found that this Euclidean distance was positively associated with tract S-A range (*r_s_*=0.688, *p*_perm_<0.001, 10,000 iterations, two-sided). This indicated that tracts forming short range connections tend to link regions with similar ranks within the cortical hierarchy, whereas longer tracts bridge across sensorimotor and association regions in the cortical hierarchy (**Figure 2d**).

**Figure 2.**
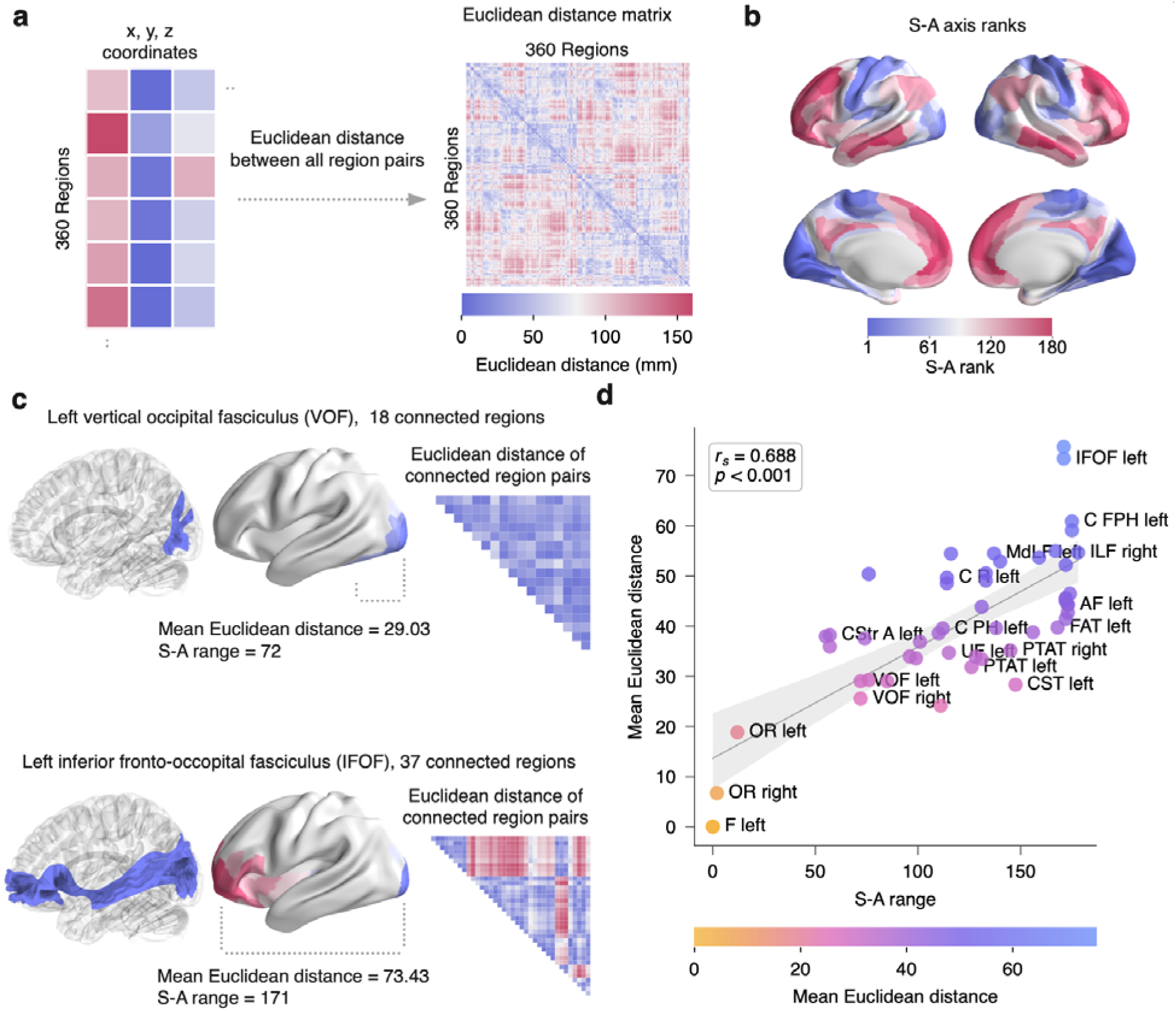
Tracts are spatially positioned to connect varying positions along the cortical hierarchy. **a)** The centroid coordinates of brain regions were used to calculate the Euclidean distance between region pairs, creating a region-by-region matrix. **b)** Cortical hierarchy was evaluated using the S-A axis, where each region is assigned a rank ranging from 1-180 per hemisphere. Sensorimotor regions have low ranks, while association regions have high ranks. **c)** Tract mean Euclidean distance was calculated from the upper triangle of the inter-regional distance matrix, using all pairs of regions connected to the tract. Tract hierarchical span, or S-A range, was defined as the maximum minus minimum S-A rank of connected regions. For example, the vertical occipital fasciculus terminates on proximal sensorimotor regions, resulting in a small S-A range and short mean distance. In contrast, the inferior fronto-occipital fasciculus connects distant frontal and occipital regions with very different hierarchical ranks, producing a large S-A range and long mean distance. **d)** Tracts forming short range connections link regions with similar hierarchical positions, whereas tracts that span long distances bridge across the poles of the cortical hierarchy. A Spearman’s correlation between the S-A axis range and the mean Euclidean distance of *N*=52 tracts was significant (*r_s_*=0.688, *p*_perm_<0.001, two-sided) as assessed using non-parametric permutation testing with 10,000 iterations. The linear fit between these measures is shown with a 95% confidence interval. WM tract labels were placed where possible while avoiding overlap.

We conducted a series of sensitivity analyses to investigate whether these relationships were consistent across different methodological choices. First, we evaluated the importance of the distance measure used to characterize the spatial embedding of tracts by using geodesic distance instead of Euclidean distance. Rather than a straight line in 3D space, geodesic distance measures the shortest path along the cortical surface between two regions. We found a similar association using Geodesic distance (*r_s_*=0.725, *p*_perm_<0.001), indicating that the distance measure used did not impact the relationship with S-A axis range. Second, to ascertain whether tract annotations were consistent across different tract types, we separately evaluated two commonly used categories of tracts: “Projection” tracts (*N*=20) and “Association” tracts (*N*=32; **Supplementary Table 1**). Similar relationships were seen in both projection (*r_s_*=0.683, *p*_perm_=0.001) and association tracts (*r_s_*=0.557, *p*_perm_=0.001). Third, we implemented an alternative significance testing framework inspired by network-rewiring approaches in connectome research^58,59^. In this tract rewiring approach, we preserved the number of regional connections of a tract, while randomly permuting the identity of the connected regions. This process generated a degree-matched but spatially random WM system, effectively eliminating the biological organization while preserving tract connection counts. This alternative significance testing approach enabled us to determine whether random, non-biologically wired WM systems of similar size would produce similar associations between tract annotations as empirical WM tracts. Using tract rewiring nulls, we again found a significant association between the Euclidean distance of connected regions and S-A axis range (*r_s_*=0.688, *p*_rewiring_<0.001; 10,000 nulls, two-sided). Together, these results demonstrate that tracts are differentially positioned within the cortical hierarchy. While some tracts tend to connect proximal regions of similar hierarchical ranking, others extend across long distances to bridge the cortical hierarchy.

### WM tracts and cognitive functions are organized along a shared spatial axis

Having characterized the spatial and hierarchical layout of tracts, we next sought to delineate the spatial relationship between WM tracts and cognitive functions. Specifically, we used partial least squares (PLS) to evaluate whether tract-function relationships align along a latent spatial axis. PLS is a multivariate analysis technique that captures dominant patterns that covary in two different data domains^60–62^ (**Figure 3a**). In our analysis, one matrix contained cognitive data (360 regions by 125 terms), representing the extent to which cortical regions are involved in each of 125 meta-analytic terms included in the *Cognitive Atlas*^38^. The other matrix was built by aggregating the tract-to-region matrix (360 regions by 26 bilateral tracts) representing tract connection probabilities across the whole brain.

**Figure 3.**
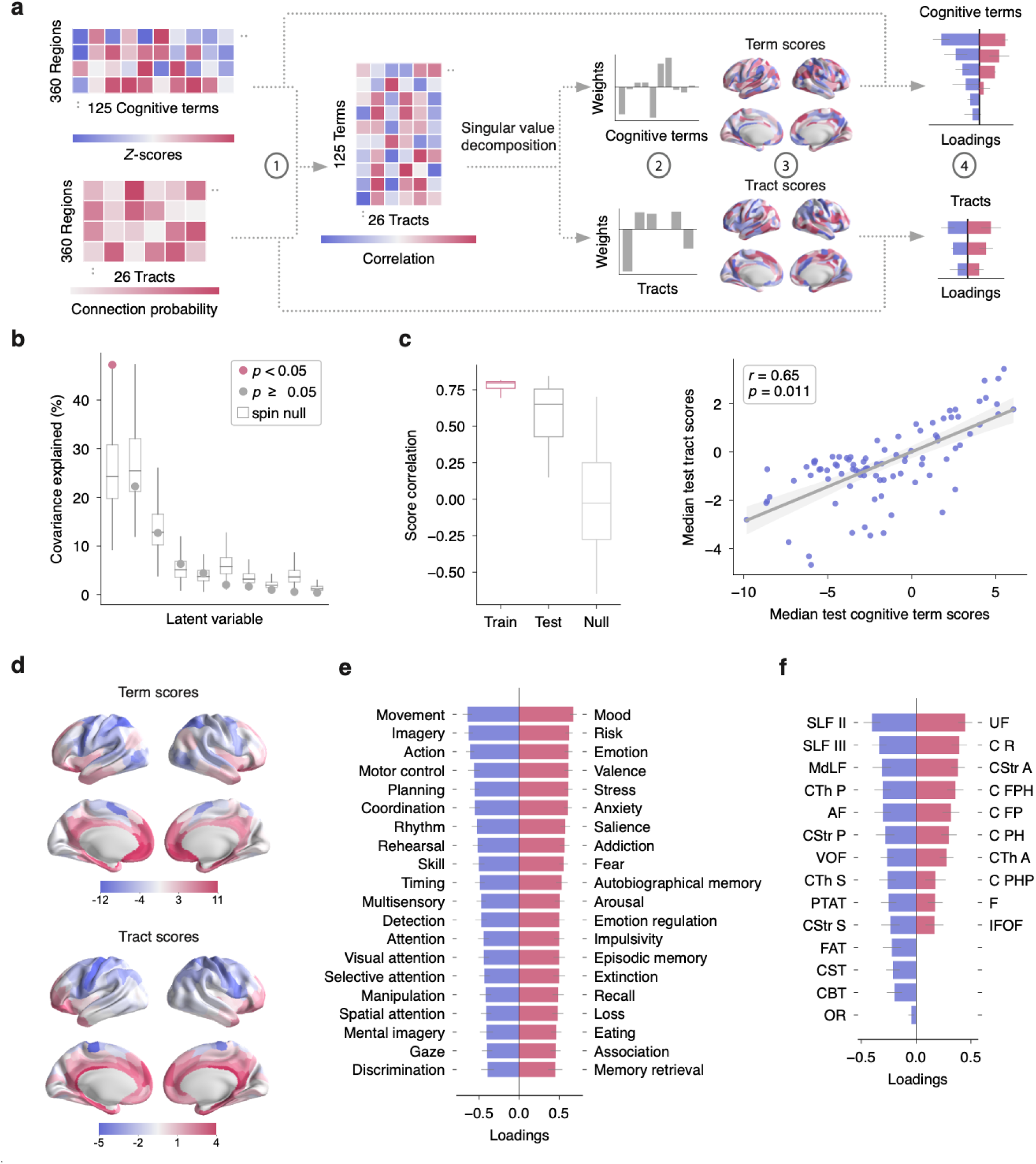
Cognitive functions and white matter tracts are spatially aligned. **a)** Diagram of steps involved in the PLS analysis. **b)** PLS identified one significant latent variable explaining 47.2% of covariance (*p*_spin_=0.002, 10,000 iterations, one-sided). **c)** The median out-of-sample correlation between term and tract scores was significant against a spatially constrained spin-based null (*r=*0.65, *p*_spin_=0.011; 1,000 repetitions, null model shown in light gray). The scatterplot shows the median test correlation of term and tract scores. **d)** The spatial pattern of cognitive term and tract scores is displayed for the first latent variable. **e)** Bars display the first latent variable loadings for the top 40 cognitive term loadings. Negative loadings (blue) and positive loadings (red) indicate opposing patterns of covariance. Error bars represent 95% confidence intervals derived from 10,000 bootstrap samples. Only reliable loadings are shown. **f)** Bars display reliable loadings for WM tracts, with similar coloring and error bar calculation as in panel e).

This approach revealed a significant latent variable that explained 47.2% of covariance (*p*_spin_=0.002, 10,000 iterations, one-sided) (**Figure 3b**). Generalizability of the PLS findings was evaluated by cross-validating the correlation between cognitive term and tract scores. Specifically, we opted for a distance-dependent cross-validation that accounts for spatial autocorrelation, in which the PLS was fit on spatially proximal regions and tested on distant regions. The median out-of-sample correlation between cognitive term and tract connections (*r=*0.65) significantly exceeded a spatially constrained spin-based null distribution of median out-of-sample correlations (*p*_spin_=0.011; 1,000 repetitions; **Figure 3c**). To evaluate the spatial alignment between cognitive terms and tracts, we generated regional “brain scores” by projecting the PLS weights back onto the original data matrices, quantifying how strongly each region expresses the shared tract-function pattern identified by the first latent variable. This analysis revealed a coherent, spatially organized pattern across the cortex, with positive scores predominantly found in ventromedial regions, and negative scores in dorsolateral regions (**Figure 3d**). Cognitive term and tract loadings from the PLS revealed a clear tract-function differentiation (**Figure 3e,f**). Tracts including the uncinate fasciculus (UF), the paraolfactory segment of the cingulum (C_R), and the anterior corticostriatal tract (CStr_A) were associated with affective functions (mood, risk, emotion). On the other hand, tracts including the superior longitudinal fasciculus (SLF II, III), the middle longitudinal fasciculus (MdLF), and the posterior corticothalamic tract (CTh_P) were primarily associated with goal-directed, action-related functions (movement, imagery, action). Taken together, findings from this analysis demonstrate that cognitive functions and WM tracts spatially co-vary, revealing a common spatial axis. This result underscores that tracts are positioned to support specific types of cognition based on their spatial layout within the brain.

### Tracts are linked to distinctive cognitive profiles

To provide researchers and clinicians with a reference that systematically characterizes the functional roles of WM tracts, we next sought to more precisely characterize the specific types of cognitive functions supported by individual tracts. To achieve this, we leveraged the cognitive terms matrix of 360 regions by 125 cognitive terms *z*-scores, reflecting the involvement of regions in each of 125 meta-analytic terms included in the *Cognitive Atlas*^38^. Next, term contributions were averaged across regions connected to the tract, resulting in 125 term contributions per tract. **Figure 4a** illustrates the term contributions for the left arcuate fasciculus. To further summarize the broader cognitive categories that tracts support, the 125 cognitive terms were grouped into the 11 categories from the *Cognitive Atlas*^38^ (**Figure 4b; Supplementary Table 2** contains a mapping of cognitive terms to each category).

**Figure 4.**
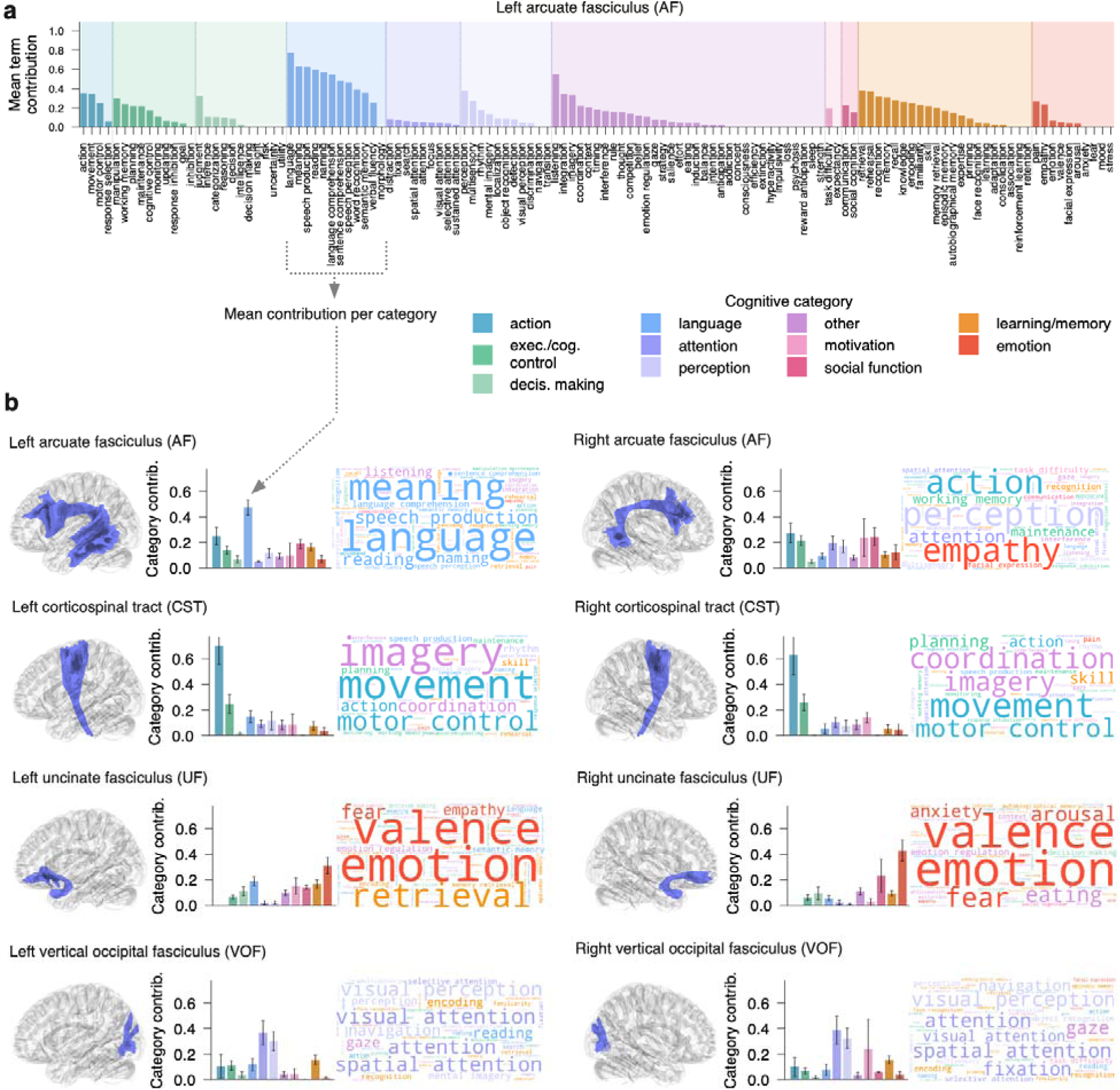
The cognitive profiles of white matter tracts. **a)** An example of cognitive term contributions in the left arcuate fasciculus (AF). *Z*-scores for each cognitive term were first normalized to the unit interval (i.e., a 0–1 scale) using robust sigmoid transformation to reduce the influence of extreme values, then masked at *z* > 1.64 (based on values in the original matrix) to retain only significant term associations. Next, term contributions were averaged across regions connected to the tract. To determine which broad cognitive categories specific tracts support, the 125 cognitive terms were further grouped into the 11 categories from the *Cognitive Atlas* and mean category contributions were calculated for each tract (shown in panel b). **b)** The contribution of each cognitive category is displayed for eight exemplar WM tracts. Each individual tract is visualized in a glass brain. Barplots indicate the contribution (mean normalized *z*-scores) of each cognitive category for the tract. Error bars reflect the standard error of the mean. Word clouds show the relative contribution of each cognitive term, colored by cognitive category. Larger words reflect higher contributions.

This approach delineated the cognitive functions associated with each tract, highlighting both expected and less well-known functions. **Figure 4b** describes the cognitive profiles of four exemplar tracts in each hemisphere (see **Supplementary Figure 1** for other tracts). For instance, the left arcuate fasciculus showed a predominant support for language functions, while also being involved in all other categories. In contrast, the right arcuate fasciculus did not show enrichment for language, with widespread involvement in all categories. The cortical spinal tract showed a clear involvement for motor control and movement, but also was linked to imagery and coordination. The uncinate fasciculus showed a predominance of emotion-related functions. Finally, the vertical occipital fasciculus supported attention and perception-related functions. Together, these findings suggest that WM tracts each have distinctive cognitive profiles, varying in both the nature and range of cognitive functions they support.

### Tracts that span the cortical hierarchy are functionally diverse

Having established the cognitive profiles of WM tracts, we next aimed to quantify the extent to which tracts are functionally specialized—supporting only a subset of cognitive functions—or functionally diverse, supporting a broad range of cognitive processes. We quantified tract cognitive diversity using the Gini coefficient, which measures how evenly its functional associations are distributed across the 125 cognitive terms. Higher values indicate a higher degree of cognitive specialization (less even distribution) and lower values indicate that these tracts support a diverse array of cognitive functions (more even distribution). Thus, the Gini coefficient provided a continuous index capturing the degree to which each tract was cognitively specialized or diverse. Specifically, we used the normalized mean term contributions of each tract (**Figure 5a**) to generate Lorenz curves representing the cumulative proportion of cognitive terms against the cumulative proportion of contributions (**Figure 5b**). Tracts closer to the equality line are relatively evenly spread across terms (e.g., the left inferior fronto-occipital fasciculus), whereas tracts that are further away from the equality line have contributions in only a small subset of terms (e.g., the left vertical occipital fasciculus). The Lorenz curves were in turn used to compute a Gini coefficient for each tract, quantifying its functional diversity (**Figure 5c; Supplementary Figure 2** contains an illustration of Gini coefficients in each of all 52 tracts separately).

**Figure 5.**
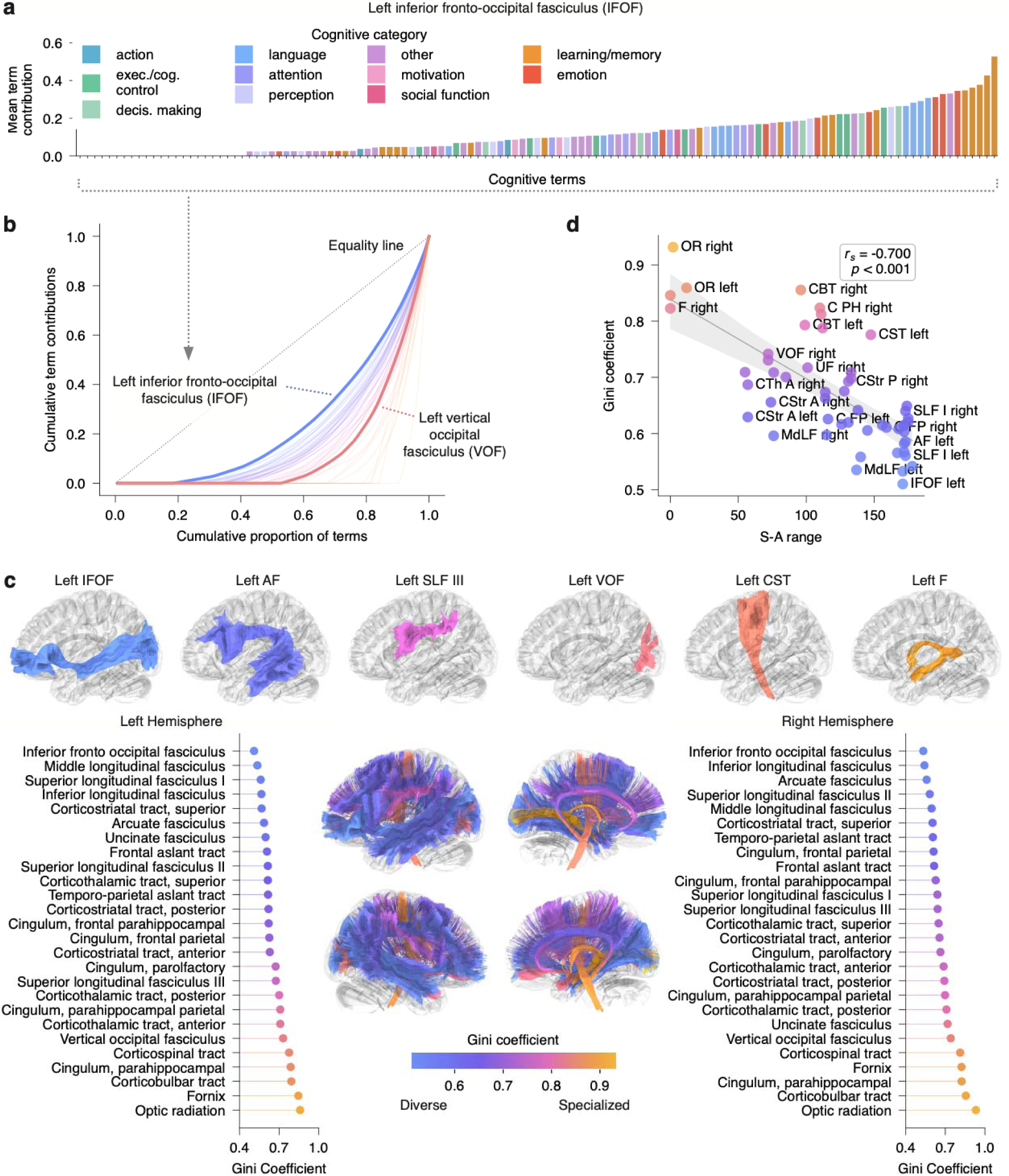
Variability in the functional diversity of white matter tracts. **a)** Term contributions were sorted in ascending order (shown for an example tract, the IFOF). **b)** Sorted terms were used to generate Lorenz curves, where the x-axis shows the cumulative proportion of term and the y-axis shows the cumulative term contributions. Each curve represents one tract; curves near the equality line indicate more diverse functional profiles (blue), while those further away are more specialized (yellow). Two example tracts are highlighted. **c)** Lorenz curves were used to obtain the Gini coefficient, quantifying the functional diversity of tracts. Low Gini values reflect tracts where most term participate equally; high Gini indicates that a few terms dominate. **d)** Tracts connecting hierarchically proximal regions were specialized, while tracts that span the cortical hierarchy were functionally diverse, as shown by a significant correlation between tract S-A range and Gini coefficient (*r_s_*=-0.7, *p*_perm_<0.001, 10,000 iterations). Tract (*N*=52) data points are colored by the Gini coefficient. The linear fit between these measures is shown with a 95% confidence interval.

This approach revealed that tracts with the greatest cognitive specialization (high Gini) included the optic radiation, the fornix, and the corticobulbar tract. In contrast, the most cognitively diverse tracts (low Gini) included the inferior fronto-occipital, the middle longitudinal, and the inferior longitudinal fasciculi. Having quantified the functional diversity of tracts, we next hypothesized that their Gini coefficients of diversity are related to embedding in the cortical hierarchy defined by the S-A axis. To test this, we evaluated the association between tract S-A range and Gini coefficient. We found a significant negative correlation (*r_s_*=-0.700, *p*_perm_<0.001), indicating that tracts connecting regions with a similar position in the cortical hierarchy are specialized, while tracts that span the cortical hierarchy support diverse cognitive functions.

Sensitivity analyses confirmed that similar relationships were seen separately in projection (*r_s_*=-0.648, *p*_perm_=0.002) and association tracts (*r_s_*=-0.597, *p*_perm_<0.001). Similarly, convergent results were found when using nulls generated with tract rewiring (*r_s_*=-0.7, *p*_rewiring_=0.01, 10,000 nulls, two-sided). Together, these results emphasize that tracts that span the cortical hierarchy are functionally more diverse than tracts that do not.

### Tract hierarchy and cognitive diversity aligns with the biological heterogeneity of cortical endpoints

To further contextualize how tracts support cognition, we examined the biological properties of regions connected to each tract. Prior work has found that structural connectivity defined using inter-regional tractography preferentially occurs between regions with similar biological profiles^20,21,31–33^; here we asked whether this would apply in long-range, anatomically-defined tracts. To do this, we selected 28 cortical properties from *Neuromaps*^8^ and *BigBrain*^12^ that describe cortical laminar cell density, myelination, and receptor density defined using Positron Emission Tomography (PET). Next, we generated a region-by-region similarity matrix containing pairwise correlations of biological properties across all regions (**Figure 6a**). Using the upper triangle of this biological similarity matrix, we calculated the mean similarity of connected region pairs for each tract (**Figure 6b**). This procedure revealed that tracts differed in the similarity of the cortical regions they connect. For example, the vertical occipital fasciculus linked regions with homogeneous biological profiles—which was reflected in a high mean biological feature similarity. In contrast, tracts like the inferior fronto-occipital fasciculus connected regions with heterogeneous biological properties, resulting in lower similarity values.

**Figure 6.**
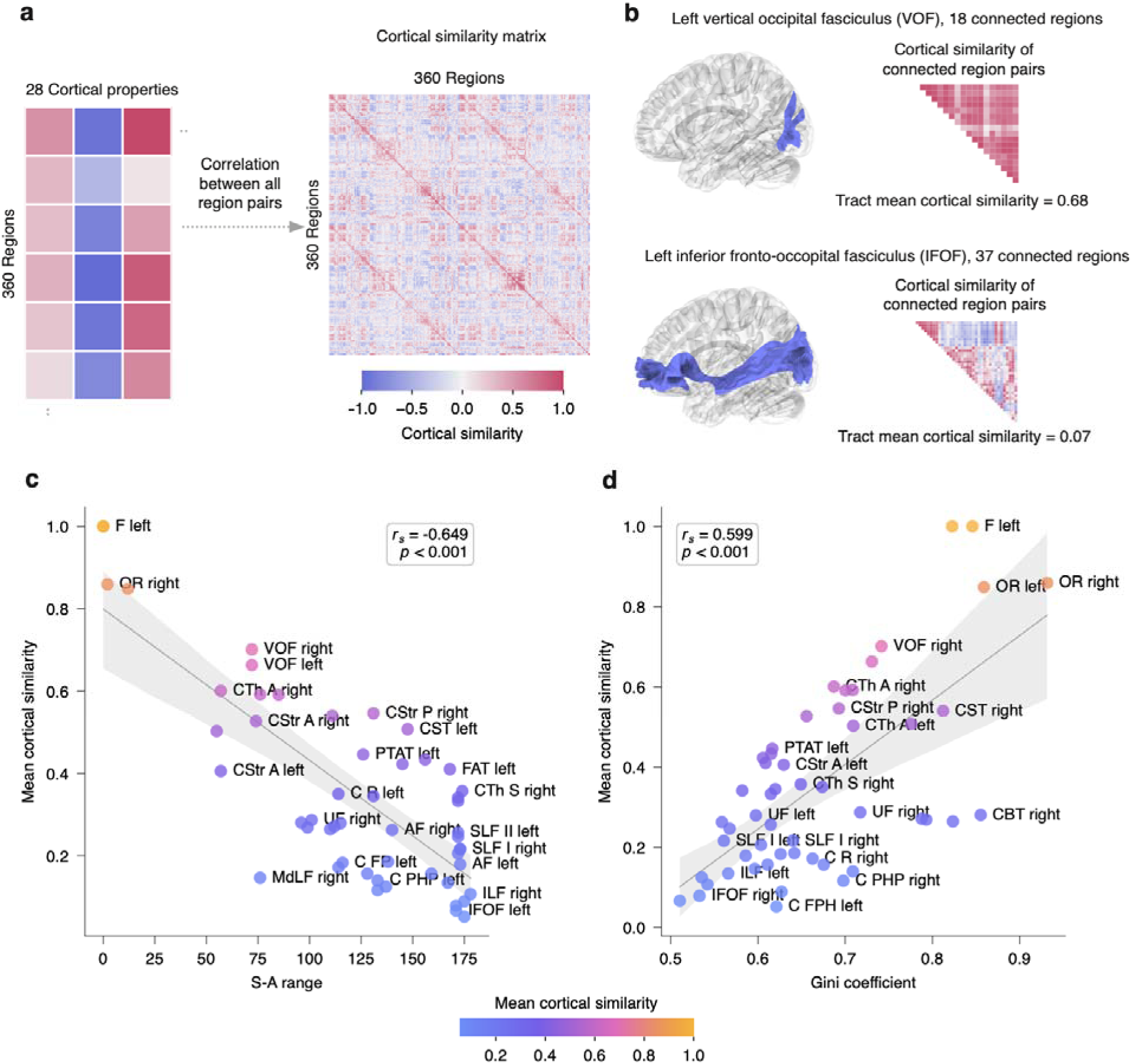
Cortical endpoint biology reflects tract hierarchy and cognitive diversity. **a)** The cortical similarity matrix was first generated as Pearson’s correlations across the 28 included cortical properties for each pair of 360 regions. **b)** Tract mean cortical similarity was computed using cortical similarity values in all region pairs connected to the tract in the upper triangle of the matrix. The vertical occipital fasciculus terminated on regions with homogeneous profile of cortical properties, a indicated by the high similarity values in all its connected regions. In contrast, the inferior fronto-occipital fasciculus connected regions with heterogeneous cortical properties, resulting in lower similarity values. **c)** Tracts bridging the cortical hierarchy had more heterogeneous biological profiles, as indicated by a significant negative correlation between tract S-A range and tract mean cortical similarity (Spearman’s *r_s_*=-0.649, *p*_perm_<0.001, 10,000 iterations). **d)** Functionally diverse tracts connected biologically heterogeneous regions, as shown by a positive Spearman’s correlation (*r_s_*=-0.599, *p*_perm_<0.001, 10,000 iterations). In both panels c) and d), the linear fit between these measures is shown with a 95% confidence interval.

Given such variation in the biological profiles of regions that were connected by tracts, we next sought to evaluate the relationship of tract biological feature similarity with a tract’s position in the cortical hierarchy (S-A range) and its cognitive diversity (Gini). We found that tract S-A axis range was negatively correlated with tract mean cortical similarity (*r_s_*=-0.649, *p*_perm_<0.001), indicating that tracts spanning the cortical hierarchy connect regions with more heterogeneous biological properties (**Figure 6c**). Furthermore, tract Gini coefficient of functional diversity was positively correlated with tract mean cortical similarity (*r_s_*=-0.599, *p*_perm_<0.001), indicating that functionally diverse tracts connect biologically heterogeneous regions.

The relationship between tract S-A range and mean cortical similarity was separately confirmed in projection tracts (*r_s_*=-0.762, *p*_perm_<0.001) and association tracts (*r_s_*=-0.462, *p*_perm_=0.008). Similar results were also seen using nulls generated with tract rewiring (*r_s_*=-0.649, *p*_rewiring_<0.001, 10,000 nulls, two-sided). We conducted the same sensitivity analyses for the relationship between tract Gini coefficients and tract mean cortical similarity and confirmed similar results (projection tracts: *r_s_*=0.610, *p*_perm_=0.006; association tracts: *r_s_*=-0.360, *p*_perm_=0.04; tract rewiring null: *r_s_*=0.599, *p*_perm_<0.001).

We additionally evaluated the choice of the measure used to characterize a tract’s hierarchical positioning by comparing tract S-A range and mean S-A—defined as the average S-A rank across connected regions in a tract—across all analyses (**Supplementary Figure 3**). This sensitivity analysis revealed that the two metrics capture different aspects of hierarchical positioning, such that compared to S-A mean, S-A range was consistently more strongly correlated with other tract properties of cortical organization, including mean Euclidean distance, Gini coefficient of diversity, and mean cortical similarity. This result indicates that tract hierarchical span is an informative descriptor of tract positioning within cortical organization.

Together, these findings suggest that tracts linking adjacent levels of the cortical hierarchy tend to connect regions with similar biological properties, whereas tracts crossing the cortical hierarchy connect regions with more heterogeneous properties. These biologically diverse connections support greater functional diversity, suggesting that these long-range tracts serve as anatomical bridges across distinct cortical environments to facilitate complex cognition.

### Tracts that span the cortical hierarchy show protracted development and are linked to individual differences in executive function

The above results showed that WM tracts are specifically positioned in the cortical hierarchy and linked to cognitive function at the population level. As a final step, we asked whether these tract features—hierarchical position and cognitive diversity—can explain variation in development and individual differences in cognition. To do this, we used data from the Philadelphia Neurodevelopmental Cohort (PNC, *N*=1,145, ages 8-23 years)^37^ to model age- and cognition-related associations in tract microstructure using generalized additive models (GAMs). A subset of 32 tracts were available for this analysis (details in Methods). Specifically, we evaluated executive efficiency, a composite factor score that captures the balance between speed and accuracy across tasks requiring executive function (including attention and working memory)^63^. We focused on executive function rather than other domains of cognition as it is known to rely on a spatially distributed network of regions. We predicted that individual variation in such executive efficiency would be linked to individual differences in long-range WM tracts that span the hierarchy^64,65^. For each WM tract, we calculated the mean fractional anisotropy (FA) representing the degree to which diffusion occurs predominantly in one direction, given that FA has been shown to track with development and cognition^66–68^. We fit tract-wise GAMs relating mean FA to age (modeled as a smooth term), adjusting for sex and motion, and with executive efficiency added for cognitive models. We quantified the effect sizes using signed partial R^2^ values.

The mean FA of tracts significantly changed with age in 30 tracts (93.8% of tracts using a false discovery rate-adjusted *p*_FDR_<0.05), with a magnitude of partial R^2^ effects in significant tracts ranging from −0.009 to 0.321; all but one tract showed significant positive associations with age. Mean FA was also significantly associated with executive efficiency in 22 tracts (68.8% of tracts at *p*_FDR_<0.05). All significant partial R^2^ effects were positive, ranging from 0.004 to 0.023, indicating that increasing tract FA was positively associated with executive efficiency.

Having described the associations between tract FA, age, and executive efficiency, we next assessed the relationship of these effects with tract annotations of hierarchical span and cognitive diversity. We found that the effect size of developmental associations were positively correlated with tract S-A axis range (*r_s_*=0.708, *p*_perm-FDR_<0.001; **Figure 7a**) and negatively correlated with tract Gini coefficient (*r_s_*=-0.445, *p*_perm-FDR_0.016; **Figure 7b**). Furthermore, executive efficiency effects were positively associated with the S-A axis range of tracts (*r_s_*=0.495, *p*_perm-FDR_=0.009; **Figure 7c**) and negatively associated with tract Gini coefficient (*r_s_*=-0.377, *p*_perm-FDR_=0.037; **Figure 7d**). These results demonstrate that tracts that span the cortical hierarchy and support greater cognitive diversity show the most pronounced developmental changes during childhood and adolescence. Furthermore, these same tracts are most strongly linked to individual variation in executive function.

**Figure 7.**
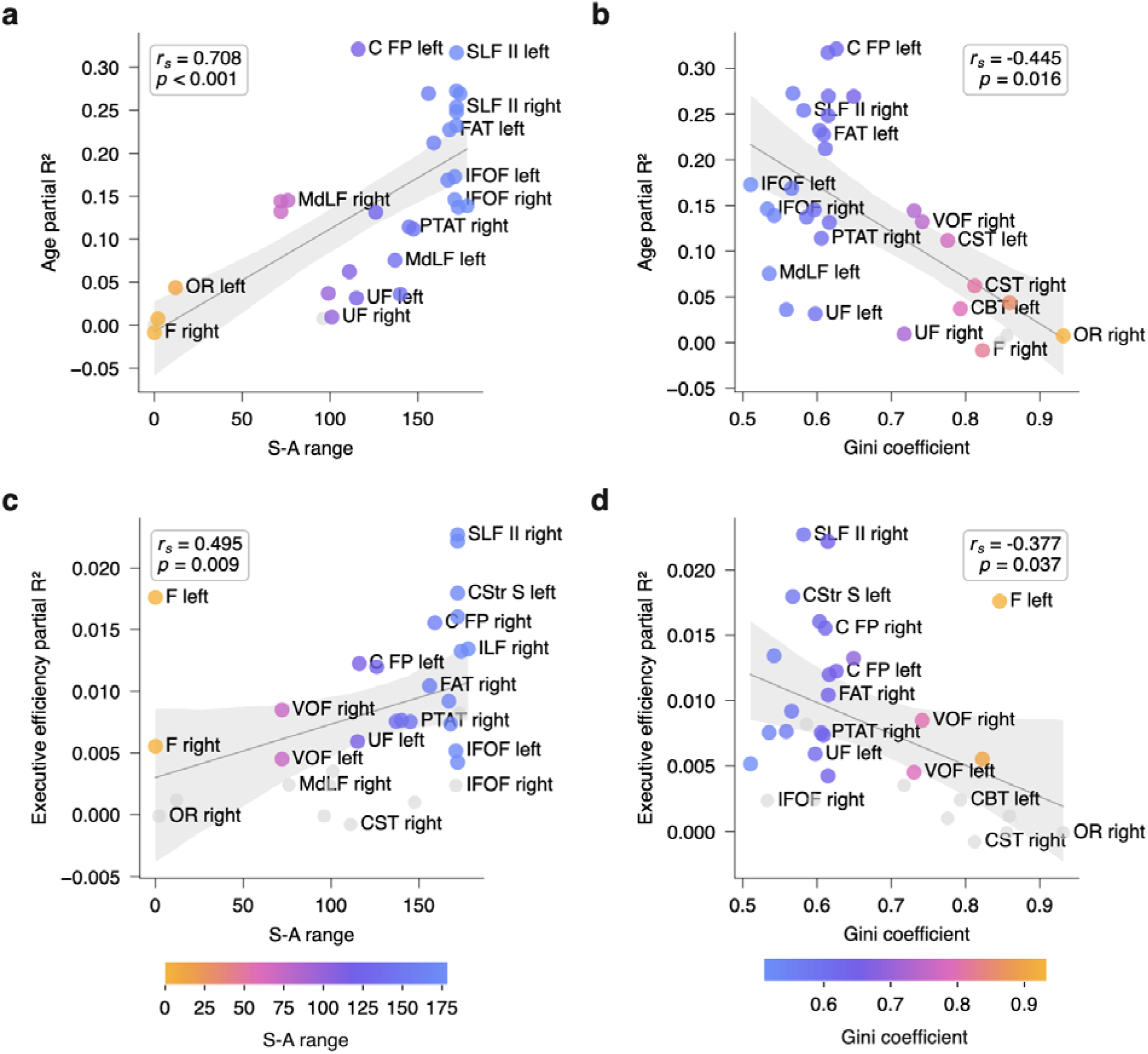
Cognitively diverse tracts that span the cortical hierarchy exhibit more development in youth and stronger links to individual differences in executive function. Individual-level analyses in the PNC (*N*=1,145, ages 8-23 years) assessed how tracts’ fractional anisotropy was associated with age and executive efficiency controlling for age (*N*=32 tracts). Effect sizes are reported as the partial R^2^ from GAMs. Tracts with non-significant partial R^2^ are plotted in gray. **a)** Tracts that spanned the cortical hierarchy developed most during childhood and adolescence, a indicated by a positive association between tract S-A range and age effects (Spearman’s *r_s_*=0.708, *p*_perm-FDR_<0.001). **b)** Age effects were also greater in magnitude for cognitively diverse tracts (low Gini), as the effect of age was negatively correlated with the Gini coefficient of tract cognitive diversity (*r_s_*=-0.445, *p*_perm-FDR_=0.016). **c)** Associations with executive function efficiency effects were more pronounced in tracts that span the cortical hierarchy (S-A range) (*r_s_*=0.495, *p*_perm-FDR_=0.009). **d)** Executive function efficiency effects were more pronounced in cognitively diverse tracts (lower Gini coefficient; *r_s_*=-0.377, *p*_perm-FDR_=0.037). Non-parametric permutation tests with 10,000 iterations were used to test the significance of the correlations and were FDR-corrected for multiple comparisons. The linear fits are shown with a 95% confidence interval.

## Discussion

Cognition does not solely arise from local activity among regions on the cortical surface – it also relies on anatomical WM tracts to coordinate neural activity across distant regions^1,2^. Here, we provide a unified framework linking anatomical WM tracts to the cognitive and neurobiological properties of their cortical endpoints. We found that tracts are differentially positioned in the cortical hierarchy to support specific cognitive functions, forming anatomical bridges to connect distinct biological environments. Notably, the position of each tract in the cortical hierarchy captures meaningful variation in brain development and individual differences in cognition. Together, these results highlight the importance of understanding WM tracts in the context of hierarchical cortical organization.

A central finding of the current study is that long-range anatomical WM tracts vary systematically in how they interface with the cortical hierarchy defined by the S-A axis^3–7^. Far from being randomly wired, canonical WM tracts are differentially positioned: some link nearby, hierarchically similar regions, while others traverse the hierarchy to connect primary sensorimotor regions with higher-order association areas. We show that such hierarchical positioning is not only intertwined with the cognitive roles of WM tracts and with the biological environments they connect, but also helped to explain developmental variation in tract microstructure and individual differences in cognition.

A majority of prior work has studied the link between brain structure and function using inter-regional connectomes. However, the inter-regional tractography that underlies structural connectomes suffers from high rates of false positives (often >50%)^26^. Furthermore, inter-regional structural connectomes are anatomically anonymous—when brain regions are found to be connected, one cannot easily ascertain which WM tract might be connecting them. Efforts to address this issue remain nascent (e.g., delineating general network properties such as efficiency^69^, or restricted to a subset of regions of interest^70^). The current study overcomes such obstacles by leveraging the tract-to-region connectome, which provides both tract identity and cortical endpoints for a set of 52 anatomical WM tracts and 360 cortical regions^35^. Importantly, the tract-to-region connectome was derived using a validated tractography approach that performs trajectory-based matching to an atlas of WM tracts curated by expert anatomists^69,71,72^. This approach effectively filters false positive connections, retaining only those matching canonical WM tracts. We leveraged this tract-to-region connectome to systematically delineate how anatomically defined WM tracts are linked to the cognitive and neurobiological properties of the cortex.

The theory that long-range WM tracts are crucial for cognition is longstanding, rooted in observations from neurological disorders caused by lesions to WM tracts, such as aphasia, agnosia, and apraxia. These disorders have historically shaped a widely adopted view that higher-order cognition arises from distributed processing across association areas, which are linked to each other and to primary sensorimotor regions via long-range WM tracts^1^. The importance of long-range WM connections was further emphasized by consistent evidence showing that these long-range connections are rare^59,73,74^. This rarity suggests that long-range WM tracts are costly to build and maintain, and therefore should form non-arbitrary, selective connections to the cortical surface that provide key benefits, including supporting complex cognition. Efforts to delineate the functional roles of major anatomical tracts have largely relied on lesion-based studies, leading to a broad identification of cognitive domains that tracts support^27,28^. Recent innovations have also started mapping fMRI activations in grey matter from a set of tasks to WM^29,30^. Here, we directly link WM tracts with cognitive functions using population-level data, yielding a systematic account of how WM tracts support cognition. Using data-driven methods (e.g., PLS), we found that dorsolateral WM tracts were linked with action-related functions, whereas tracts with ventromedial connections were linked to affective functions. This finding supports the view that tracts are positioned to support specific types of cognition based on their spatial layout within the brain.

In addition to these broad patterns of brain organization, we used meta-analytic task activations to delineate the unique cognitive profiles of tracts. This analysis confirmed well-established functions, including language for the arcuate fasciculus^75,76^ and movement for the corticospinal tract^77^. Furthermore, we found evidence for less commonly discussed roles, such as the right arcuate fasciculus’ involvement in socio-emotional processes like empathy^78^ or the corticospinal tract’s association with mental imagery^79^. By systematically delineating the cognitive profiles of anatomical WM tracts using population-based data, this work provides a practical reference for researchers and clinicians seeking to understand the typical functions associated with specific long-range tracts.

Observing that the range of functions varied across tracts, we sought to quantify the cognitive diversity of tracts. Despite widespread assumptions that long-range WM tracts support complex and diverse cognitive processes, surprisingly few empirical efforts have examined this claim. Our findings indicate that WM tracts vary along a continuous spectrum from cognitive specialization to diversity, ranging from highly specialized tracts such as the optic radiation, the fornix, and the corticobulbar tract, to diverse tracts such as inferior fronto-occipital, the middle longitudinal, and the inferior longitudinal fasciculi. While prior work has considered cognitive diversity primarily at the regional level^80,81^, our results emphasize that the cognitive diversity of anatomical WM tracts is dictated by their positioning in the cortical hierarchy: the more a tract spans the S-A axis, the greater its cognitive diversity. This result echoes classic theories proposing that complex cognition may be enabled by structural connections linking primary sensorimotor regions to higher-order association cortex^1^, while also providing new evidence showing that tracts bridging hierarchical poles support cognitive diversity.

Importantly, cognitive functions depend on the neurobiological properties of the cortical regions supporting them. Multiple lines of evidence suggest that several of these surface properties are tightly coupled to wiring patterns in the human brain through evolution and development. Intracortical myelination and WM tracts undergo pronounced maturational changes during the same developmental window^82,83^, and recent evidence suggests that the maturation of WM structure and intracortical myelin is coupled^23^. Common evolutionary and developmental principles are thought to orchestrate the patterning of laminar thickness across the cortex and the layer-specific formation of structural connections^19,21,84,85^. Gene expression is coordinated in structurally connected regions, and these regions exhibit an enrichment for genes involved in brain development, when structural connections are formed^18^. Similarly, neurotransmitter receptor distribution closely aligns with the bain’s structural connectivity^15^. Despite such evidence suggesting that regional biological properties are related to WM architecture, it remained unknown how tracts were positioned within the neurobiological milieu of the cortical surface. Evidence from some animal models supported the idea that structural connections were preferentially made between regions with similar neurobiological profiles – even when those regions were distant from each other^20,21,31–33^. However, others have shown that the longest structural connections in fact seemed to promote neurobiological diversity by connecting disparate regions^34^. By examining the neurobiological similarity of regions connected to long-range anatomically-defined tracts, we observed that their biological profiles vary systematically with their positioning along the cortical hierarchy. Specifically, tracts that connected regions with a similar position on the S-A axis had greater neurobiological homogeneity, whereas tracts that spanned the hierarchy tended to maximize neurobiological diversity. These results may thus largely resolve the discrepancy of existing literature, suggesting that only some WM tracts preferentially connect biologically similar regions, while others bridge biologically heterogeneous environments to support cognitive diversity.

Situating WM tracts in the cortical hierarchy also explained variation in tract development and individual differences in cognition. Numerous studies have reported dramatic changes in WM microstructure during childhood and adolescence, with variation in the magnitude and timing of these changes depending on the tracts^86–89^. Abundant evidence also suggests links between tract microstructure and individual differences in cognition^90,91^. By convention, WM tracts are typically separated into tract types, including projection, association, and commissural tracts. While the magnitude of age and cognition differences appear to vary by tract type^86^, such broad subdivisions only provide a coarse understanding of WM maturation and its relevance for cognitive performance. Here, our results provide a more fine-grained, continuous characterization of WM tracts that is rooted in the hierarchical organization of the cortex. We found that age and cognition effects were more prominent in hierarchy-spanning WM tracts that supported greater cognitive diversity.

Why might tracts with wide hierarchical spans supporting diverse functions show greater associations with development and cognition? While animal studies are needed to elucidate the mechanisms that link developmental sequences of WM to tract positioning within the cortical hierarchy, a wealth of evidence documents the extensive cortical refinement that happens in association cortices during adolescence^92,93^. Notably, this cortical remodeling has been suggested to drive activity-dependent changes in WM^94,95^. We speculate that activity-driven developmental changes in association regions that undergo protracted development similarly drives greater WM remodeling in connected tracts. In contrast, tracts that connect primary sensorimotor regions have likely already reached maturity by late childhood. Tracts with a wider hierarchical span may show the most pronounced age-related changes in adolescence because they must integrate information between primary and association territories. This is consistent with recent work in humans showing systematic developmental variation along the length of WM tracts, with tract endpoints near sensorimotor tract endpoints maturing earlier than endpoints in higher-ranking association regions^89^. Together, these findings suggest that long-range WM tracts maturation may unfold through hierarchical cascades, shaped by interactions with a heterochronous program of cortical maturation that progresses from lower order sensorimotor regions to higher-order association cortices.

Our findings should be considered in the light of several limitations. First, this study used publicly available data and was therefore constrained in the tractography method and tract types that were examined. Second, this study examined population-level trends in tract-to-cortex relationships. While most connection probabilities were close to either 0 or 1, it is important to note that there is some inter-individual variability in tract connection patterns^35^; understanding this variability remains an important area for future research. Third, due to changes in the *DSI Studio* WM atlas over time, a subset of 32 tracts out of the 52 tracts was used to quantify development and cognition effects in tract microstructure. Fourth, we examined individual differences in tract microstructure using fractional anisotropy (FA), which indexes the degree to which water diffusion is directionally constrained. FA is influenced by multiple factors, including crossing fibers, fiber diameter and density, and myelination^96^.

Taken together, our findings provide not only a better understanding of how WM is situated within the context of cortical organization, but also suggest a framework for mapping individual differences in WM development and cognition. In doing so, this work emphasizes that understanding the role of WM tracts requires moving beyond conventional categories of projection, association, and commissural tracts and toward a cortex-anchored understanding of how anatomical WM tracts support cognition. This study opens several avenues for future research at the intersection of WM architecture, cortical organization, and cognition. Replication and extension using atlases that include commissural and subcortical tracts will help refine the current framework. The measures of tract distance, hierarchy, cognitive diversity, and biological milieu generated here can also be leveraged to interpret other individual-level WM differences, including the impact of childhood environment and aging. Moving forward, understanding how evolution and cortical expansion gave rise to such a wide range of WM tract connection patterns across the cortical hierarchy, and how these patterns relate to the emergence of higher-order cognition, represents an important direction for future research. Longitudinal work in developmental datasets will be invaluable to investigate the lagged relationships between cortical function and the development of WM tracts. Finally, understanding how interindividual variability in tract-to-region connections relates to development, cognition, cortical hierarchy, and known variability in person-specific functional networks^97,98^ is an essentially untapped domain for future discoveries in personalized neuroscience.

## Methods

### Population-level data

#### Tract-to-region atlas

Probabilistic tract-to-region connections were obtained from a publicly available population-level atlas^35^. This atlas was created by reconstructing 52 tracts in 1,065 young adults from the Human Connectome Project^99^ (age range: 22-37 years, mean age: 28.75 years) with *DSI Studio* (http://dsi-studio.labsolver.org)^100^. In each individual, 52 WM tracts were reconstructed based on multishell diffusion scans (the consortium paper contains detailed information on the acquisition scheme^99^) using a combination of deterministic fiber tracking, randomized parameter saturation, topology-informed pruning, and trajectory-based recognition (see Ref.^35^ for details). Next, each tract’s connections to all 360 cortical regions of the Human Connectome Project’s multimodal parcellations (HCP-MMP) atlas^9^ were evaluated in each individual, producing a binary (absence vs. presence) tract-to-region mapping per participant. A connection was considered present when tract voxels intersected with the cortical region in volumetric space. A population-level tract-to-region matrix was then defined as the proportion of individuals with a given tract-to-region connection^35^. This proportion, termed “connection probability”, ranged from 0, indicating that no participant exhibited a given tract-to-region connection, to 1, reflecting that all participants had the connection. We used a connection probability threshold of 0.5 to establish the presence of a tract-to-region connection in all analyses. Anatomical tracts considered for the tract-to-region connections included projection and association tracts; cerebellar, commissural, brainstem, or connections between subcortical structures were excluded due to methodological limitations (see Ref.^35^ for details). The resulting tract-to-region connectome is a matrix of *m* brain regions by *n* WM tracts (i.e., a 360-by-52 matrix), where each cell corresponds to the population probability of a WM tract structurally connecting a surface brain region. Note that because this analysis focused on within-hemisphere tracts, the full matrix contained NAs for the contralateral regions of each tract. Critically, this novel form of tract-to-region connectome provides information about which WM tracts are involved and where they connect to the cortex, offering a biologically grounded alternative to more common region-to-region structural connectomes that lack tract identity information and are prone to high rates of false positives^26^.

#### Meta-analytic Cognitive Atlas terms

Cognitive term maps were obtained from *Neurosynth*^36^ version 0.7, a meta-analytic tool that synthesizes task functional MRI results from over 15,000 published studies by searching for associations between MRI voxel coordinates and frequently mentioned keywords in the abstracts of the articles. Values in these cognitive term maps are association test *z*-scores quantifying to what extent a given cortical region was more consistently activated in studies that mentioned the term, compared to studies that did not mention it. For example, large positive *z*-scores in the motor cortex for the “action” term indicates that studies reporting “action” in their abstract were more likely to report activations in that region compared to studies that did not mention it. In line with prior work^101^, we drew on the *Cognitive Atlas*^38^ (http://www.cognitiveatlas.org/) to select terms within the *Neurosynth* database that correspond to well-characterized cognitive functions reflecting the current consensus of cognitive neuroscience. This process resulted in a set of 125 cognitive terms that ranged from lower- to higher-order cognitive processes. **Supplementary Table 2** contains a full list of the cognitive terms used. The Neuroimaging Meta-Analysis Research Environment^102^ (*NiMARE*) was used to derive meta-analytic task activation maps for these 125 terms in volumetric space using Multilevel kernel density (MKDA) Chi-square analysis^103^. We used *Neuromaps*^8^ to map volumetric meta-analytic statistical maps to *fsLR32k* surface space and parcellate them into HCP-MMP atlas^9^ regions. *Neuromaps* is a toolbox that provides a number of native space maps as well as functionality for converting to different spaces and resolution to facilitate across-map comparisons.

#### Biological cortical features

Biological features were selected based on their conceptualized relevance to WM anatomy. First, we selected T1w/T2w maps as proxies of intracortical myelination^104,105^ as developmental changes in intracortical myelination have been observed alongside changes in anatomical WM tracts^82,83^, and recent evidence suggests that the maturation of WM structure and intracortical myelin is coupled^23^. Second, previous reports suggest that gene expression is coordinated in regions that share structural connections, particularly genes relevant for neuronal development and axon guidance^18^. Therefore, we selected the first principal component of gene expression^10,11^, representing dominant patterns of variation across cortical regions. Moreover, due to the relationship between WM structure and neurotransmission^15^, we included a synapse density map (measured via PET tracer binding to the synaptic vesicle glycoprotein 2A (SV2A)^45,49,57^, as well as 19 receptors and transporters spanning nine neurotransmitter systems, including serotonin (5-HT_1A_, 5-HT_1B_, 5-HT_2A_, 5-HT_4_, 5-HT_6_, 5-HTT)^106–108^, histamine (H_3_)^15,109^, dopamine (D_1_, D_2_, DAT)^15,110–112^, norepinephrine (NET)^15,113^, acetylcholine (α_4_β_2_, M_1_, VAChT)^15,114–116^, cannabinoid (CB_1_)^117^, opioid (MOR)^118^, glutamate (mGluR_5_, NMDA)^15,119–122^, and GABA (GABA_A/BZ_)^123^. All neurotransmitter maps were measured from positron emission tomography (PET). Third, we included laminar thickness for six cortical layers (I-VI), given that laminar organization and long-range WM tracts are thought to emerge through common ontogenetic principles^19,21,84,85^.

Biological cortical features including intracortical myelination, gene expression, and neurotransmitter density maps, were obtained through *Neuromaps*^8^. Laminar thickness maps were obtained through the *BigBrainWarp* toolbox^12^, which derived these measures by applying an automatic segmentation technique to the BigBrain high-resolution (20×20×20 μm) histological atlas of a post-mortem human brain (65 year-old male)^124–126^. More information on the estimation of cortical layer thickness data is provided in Wagstyl *et al.*, 2020^14^. All 28 maps were transformed from their original space to *fsLR32k* surface space and parcellated into 360 HCP-MMP regions using *Neuromaps*^8^ tools.

### Tract annotations

Using these population-level tract-to-region, cognitive, and biological cortical maps, we computed several “tract annotations”, each describing WM tracts based on the properties of the cortical regions it connects. As described below, these annotations quantify different aspects of how a tract is positioned within cortical organization.

#### Tract mean Euclidean distance

To characterize the spatial embedding of tracts, we used the 3D coordinates of the 360 HCP-MMP regions and generated a region-to-region matrix containing the Euclidean distance for each region pair. The Euclidean distance reflects the straight-line connecting two regions in 3D space. To define a tract’s spatial embedding, we selected the upper triangle (excluding the diagonal) and subsetted region pairs connected to the tract. We then quantified each tract’s spatial embedding as the mean of Euclidean distances in connected region pairs.

#### Tract mean geodesic distance

To assess the importance of the distance measure used to characterize the spatial embedding of tracts, we conducted a sensitivity analysis using geodesic instead of Euclidean distance. The geodesic distance is the shortest path along the cortical surface between two regions. The *brainSMASH* toolbox^127^ was used in combination with *Connectome Workbench*, version 2.0.1 (https://www.humanconnectome.org/software/connectome-workbench) to calculate geodesic distances for the *fsLR32k* midthickness surface. Similar to the method described above for Euclidean distance, the upper triangle of the region-to-region geodesic distance matrix was then selected (excluding the diagonal) and region pairs connected to the tract were selected. The geodesic distance for each tract was then calculated as the mean of geodesic distances of connected region pairs. We then calculated the correlation between this tract geodesic distance annotation and the tract S-A range to determine whether the association was similar to what was observed using tract Euclidean distance.

#### Tract S-A range

To delineate the hierarchical span of tracts, we leveraged the regional S-A axis ranks identified in prior work as a measure of cortical hierarchy^3^. This axis was delineated by averaging the rank orderings of ten multimodal cortical properties that manifest systematic variation from primary sensorimotor regions to higher order association regions. The resulting cortical S-A map reflects the average hierarchical position of each region. The S-A cortical map for the HCP-MMP parcellation was obtained from https://github.com/PennLINC/S-A_ArchetypalAxis/tree/main/Glasser360_MMP. We defined a tract’s hierarchical span as the tract’s S-A range, calculated by taking the maximum minus minimum S-A rank of regions connected to the tract.

#### Tract cognitive term contributions

We used the matrix composed of 360 regions x 125 *Cognitive Atlas* terms to characterize the cognitive profiles of tracts. To reduce the influence of extreme values, *z-*scores for each cognitive term were first normalized to the unit interval (i.e., a 0–1 scale) using a non-linear scaled robust sigmoid transformation from the *abagen* Python package^11^. We then defined a mask as *z* > 1.64 in the original *z*-scores matrix and applied to the normalized matrix to retain only significant positive term associations. Next, term contributions were averaged across all regions connected to the tract. This process resulted in a terms-by-tracts matrix (125×52) representing the mean normalized contribution of each term in each tract. To further summarize the cognitive profiles of tracts, we used the 11 categories from the Cognitive Atlas (obtained at https://www.cognitiveatlas.org/concepts/categories/all; see **Supplementary Table 2** for a list of terms associated with each category) and computed the mean and standard error of the mean (SEM) of all of a tract’s term contributions per cognitive category. Categories included “Action”, “Learning and Memory”, “Emotion”, “Attention”, “Reasoning and Decision Making”, “Executive/Cognitive control”, “Social Function”, “Perception”, “Motivation”, “Language” and “other”. Each tract was thus associated with average contributions in each of the 11 cognitive categories. Contributions ranged from 0, indicating that the tract did not contribute to the cognitive category, to 1, indicating that the tract is heavily involved in the category.

#### Tract Gini coefficient

To assess the cognitive diversity of WM tracts, we computed the Gini coefficient. Commonly used in economics to measure income inequality, the Gini coefficient in this context quantifies how evenly distributed a tract’s functional associations are across the 125 cognitive terms, providing a measure of functional diversity vs. specialization. The Gini coefficient of a tract is calculated by sorting all its term contributions in ascending order and computing two measures. First, we calculated the cumulative proportion of terms, representing the proportion of the population (terms) accounted for at any given point in the distribution. Second, we calculated the cumulative term contributions, reflecting the actual term contributions at the same point in the distribution. These two measures can be visualized as Lorenz curves, where the diagonal represents the perfect equality line in which all terms participate equally. The Gini coefficient is the ratio of the area between the equality line and the Lorenz curve to the total area under the equality line. A Gini coefficient of 1 corresponds to a maximally unequal contribution pattern, where one cognitive term dominates the distribution. This represents a highly specialized tract that supports only one cognitive function. A Gini coefficient of 0 indicates that all terms contribute equally, and reflects a highly diverse tract.

#### Tract biological cortical similarity

To quantify the biological similarity of cortical features linked to each tract, we first computed a region-to-region similarity matrix based on the 28 biological cortical feature maps (see “Biological cortical properties” section). Similarly to prior work^15,128^, the regional similarity matrix was generated by computing the correlation (Pearson’s *r*) between biological profiles for each region pair. The similarity matrix thus represents the extent to which pairs of regions exhibit the same biological blueprint. To define a tract’s biological cortical similarity, we calculated the mean *r* value within the upper triangle (excluding the diagonal) of connected region pairs.

### Significance testing

#### Non-parametric permutation testing

A key goal of this study was to assess the relationships between the various tract annotations generated based on various cortical properties as described above. We evaluated these associations between tract annotations using Spearman’s rank correlations, denoted *r_s_*. Note that all tract annotations used in these analyses were derived from group-averaged population maps (e.g., probabilistic tract-to-region connections and cortical feature maps). Each annotation represents a single summary measure per WM tract (e.g., Euclidean distance, S-A range, etc.), computed based on cortical properties across the set of cortical regions connected to that tract. Thus, the associations evaluated here correspond to population-level summary measures of WM tracts, rather than subject-level variability. To assess the significance of the associations, we applied non-parametric permutation with 10,000 iterations. Each iteration randomly shuffled the Y variable while keeping the X variable fixed, and re-correlated the X and Y, thus disrupting the original relationship between both variables and generating a null distribution of correlations. The significance of the relationship was determined by calculating the proportion of permuted correlations with absolute values greater than or equal to the absolute value of the empirical correlation, providing a two-sided *p*-value (denoted *p*_perm_) that tests against the null hypothesis of no association between the variables.

### Partial Least Squares analysis

Partial Least Squares (PLS) analysis was used to investigate the relationship between population-level maps of cognitive terms and tract-to-region connection probabilities. PLS is a data-driven, multivariate approach that takes cognitive terms and tracts matrices as inputs and decomposes them into orthogonal latent variables with maximum covariance^129^.

#### Data inputs

The cognitive terms matrix included population-level activation values in 360 regions for 125 cognitive terms. The WM tracts matrix was generated by aggregating left and right tracts into a single matrix of 360 regions and 26 tracts, representing population-level tract connection probabilities across the whole brain. In this matrix, each column represented a tract bilaterally (e.g., the left and right arcuate fasciculi were aggregated into a single arcuate fasciculus column). Both the cognitive terms and the tract connection input matrices were *z*-scored prior to running PLS.

#### Latent variables and weights

Each latent variable generated by the PLS comprised cognitive term weights, tract weights, and a singular value indicating the amount of covariance explained by the latent variable. Positively weighted cognitive terms covaried with positively weighted tracts, while negatively weighted cognitive terms covaried with negatively weighted tracts.

#### Brain scores

PLS-derived vector weights were projected back onto the original data to estimate “brain scores” that represent the extent to which each brain region expresses the cognitive term weights and the tract weights identified by the latent variable. Thus, positive brain region scores indicated that the region expressed covariance between positive cognitive term weights and positive tract weights. Conversely, negative brain region scores indicated that the region expressed covariance between negative cognitive term weights and negative tract weights.

#### Cognitive term and tract loadings

Loadings for the cognitive terms and tracts were computed as the Pearson correlation between the original data matrices and the corresponding regional brain scores. Cognitive term and tract loadings can be understood as the extent to which a given cognitive term or tract contributed to the pattern identified by the latent variable. To estimate the reliability (i.e., 95% confidence intervals) of the PLS loadings, we used bootstrap resampling, where brain regions (rows) of the original X and Y data matrices were randomly selected with replacement 10,000 times. PLS was then reapplied to each resampled dataset, producing a distribution of bootstrap-resampled loadings.

#### Significance testing of the latent variables

The significance of the association between regional term and tract matrices in the PLS was evaluated using spatial autocorrelation preserving permutations (“spin tests”)^130,131^. Spin-tests were used because the data inputs were population-level whole-brain maps where rows are brain regions, not subject-level data (where rows would be individuals). In this analysis, the spin test implemented the null hypothesis that cognitive terms and tracts were correlated only due to inherent spatial autocorrelation. This implementation was done by spinning the indices of regions in the cognitive terms matrix, and recomputing the PLS on the spun terms matrix and original tracts matrix. This process was repeated for 10,000 spun versions of the cognitive terms matrix, resulting in a null distribution of the amount of covariance explained by each of the latent variables. The *p*-value (denoted *p*_spin_) represented the probability of the observed association between cognitive terms and tracts matrices purely due to comparable spatial autocorrelation, and was calculated as the proportion of null singular values that exceeded the observed singular value for each latent variable.

#### Cross-validation

In line with previous neuroimaging applications of PLS^132–134^, we evaluated the generalizability of the PLS findings by cross-validating the Pearson’s correlation between cognitive terms and tract scores in a distance-dependent manner. We opted for this approach, as it accounts for the fact that nearby regions tend to have more similar profiles, and therefore random splits may overestimate model performance due to spatial autocorrelation. By maximizing the spatial distance between regions of each test and training set, the cross-validation provides a more conservative way to estimate the generalizability of the PLS. Specifically, each training set was selected using a random “seed” region and 75% of the regions closest to it in Euclidean distance; the test set comprised the remaining 25% most distant regions. We then re-ran the PLS on each training set (75% of regions). Test sets were projected onto the singular vector weights from the corresponding training sets to generate a distribution of out-of-sample predicted term and tract scores, along with predicted correlations. The significance of the median out-of-sample correlation was determined by comparing the observed median test correlation against a null distribution generated from 1,000 spin-based permutations that preserved spatial autocorrelation while randomizing the cognitive term-tract relationships. For each spin iteration, the same distance-based cross-validation procedure was applied to rotated data and the median test correlation was recalculated, creating a null distribution of spatially-constrained median test correlations. The *p*-value was computed as the proportion of null out-of-sample median correlations that were greater than the observed median test correlation.

### Sensitivity analyses

We performed several sensitivity analyses to ascertain whether the findings were impacted by methodological choices made in the study.

#### Analyses by tract type

First, to ascertain whether tract annotations were consistent across different tract types, we divided the tracts into two types: “Projection” tracts (*N*=20) and “Association” tracts (*N*=32). **Supplementary Table 1** contains the assignments of each tract into the two types. We then recomputed the correlations between tract annotations separately in each tract type.

#### Significance testing using tract rewiring

Second, to assess the robustness of our findings across different significance testing frameworks, we implemented a tract-based null model inspired by network-rewiring approaches in connectome research^58,59^. This “tract rewiring” procedure preserved the number of region connections per tract (degree-preserving), but randomized which regions were connected, constrained to remain within the same hemisphere as the original tract. This was repeated to generate 10,000 rewired tract-to-region matrices, which all maintained the overall structural properties of the tract “network” while disrupting specific regional connectivity patterns. This allowed us to test whether a randomly wired WM system—with the same tract connection counts—would produce similar associations between tract annotations as what was observed using empirical tract-to-region connections.

### Individual-level data

#### Participants

Participants were drawn from the Philadelphia Neurodevelopmental Cohort (PNC)^37^, a cross-sectional dataset that comprises a community sample of youth from the greater Philadelphia area. A sample of *N*=1,145 participants with demographic and diffusion imaging data were available after applying inclusion criteria; of these, *N*=1,142 participants had cognitive data. The sample age range was 8-23 years (mean of 15.3±3.5 years); it included 537 males and 608 females (sex was self-reported; intersex was not assessed); race and ethnicity distributions were: *N*=3 American Indian or Alaskan Native, *N*=10 Asian, *N*=486 Black or African American, *N*=120 identifying as multiracial and *N*=525 White. All participants over the age of 18 years gave written informed consent before study participation. Participants under the age of 18 years gave informed assent with written parental consent. All individuals received monetary compensation for participation in the study. All study procedures were approved by the Institutional Review Boards of both the University of Pennsylvania and the Children’s Hospital of Philadelphia.

#### Cognitive assessment

Cognition was assessed using the Penn computerized neurocognitive battery (CNB), comprising 14 tests adapted from tasks applied in functional neuroimaging to evaluate a broad range of cognitive domains^135,136^, including executive function (abstraction and flexibility, attention, working memory), episodic memory (verbal, facial, spatial), complex cognition (verbal reasoning, nonverbal reasoning, spatial, processing), social cognition (emotion identification, emotion intensity differentiation, age differentiation), motor and sensorimotor speed. Building upon prior work that delineated the factor structure of cognitive performance in the PNC^63^, we selected executive efficiency to measure cognitive performance, as it captures the balance between speed and accuracy across tasks requiring executive function and is thought to be supported by distributed cortical regions that communicate through long-range WM tracts^64,65^.

#### MRI data acquisition

T1w structural MRI and diffusion weighted imaging (DWI) data from the PNC were used to reconstruct WM bundles from participants in the PNC. All PNC MRI data were collected on the same 3T Siemens TIM Trio Scanner and 32-channel head coil at the University of Pennsylvania for all participants. T1w images were acquired with a magnetization-prepared rapid acquisition gradient-echo (MPRAGE) sequence with the following parameters: TR = 1,810 ms, TE = 3.51 ms, TI = 1,100 ms, flip angle = 9 degrees, 160 slices, and voxel resolution = 0.94 × 0.94 × 1 mm.

DWI scans were acquired using a twice-refocused spin-echo (TRSE) single-shot echo-planar imaging (EPI) sequence with the following parameters: TR = 8,100 ms, TE = 82 ms, and voxel resolution = 1.875 × 1.875 × 2 mm. The full 71 volumes, 64-direction set was divided into two independent 32-directions imaging runs, for a total scanning time of 11 min. Each 32-direction subset was chosen to be maximally independent such that they separately sampled the surface of a sphere. The complete sequence consisted of 64 directions with b = 1000 s/mm^2^ and 7 interspersed scans with b = 0 s/mm^2^.

#### Diffusion MRI preprocessing

The T1w structural images and the DWI scans were preprocessed using *QSIPrep* 1.0.0rc1^137^, an integrative pipeline for processing diffusion-weighted MRI data that uses the software tools described below. *QSIPrep* is based on *Nipype* 1.9.1^138,139^. The text in the following sections was automatically generated in *QSIPrep* and released under the CC0 license for reuse in manuscripts.

#### Anatomical preprocessing

The anatomical reference image was reoriented into AC-PC alignment via a 6-DOF transform extracted from a full Affine registration to the MNI152NLin2009cAsym template. A full nonlinear registration to the template from AC-PC space was estimated via symmetric nonlinear registration (SyN) using *antsRegistration*. Brain extraction was performed on the T1w image using *SynthStrip*^140^ and automated segmentation was performed using *SynthSeg*^141,142^ from FreeSurfer version 7.3.1.

#### Diffusion data preprocessing

A total of 2 DWI series in the j-distortion group were concatenated, with preprocessing operations performed on individual DWI series before concatenation. Any images with a *b*-value less than 100 s/mm^2^ were treated as a *b*=0 image. MP-PCA denoising as implemented in *MRtrix3*’s *dwidenoise*^143^ was applied with an auto-voxel window. When phase data were available, this was done on complex-valued data. After MP-PCA, Gibbs unringing was performed using TORTOISE’s Gibbs^144^. Following unringing, the mean intensity of the DWI series was adjusted such that the mean intensity of the b=0 images matched across each separate DWI scanning sequence. B1 field inhomogeneity was corrected using *dwibiascorrect* from MRtrix3 with the N4 algorithm^145^ after corrected images were resampled.

*FSL* (version 6.0.7.9)’s *eddy* was used for head motion correction and Eddy current correction^146^. *Eddy* was configured with a *q*-space smoothing factor of 10, a total of 5 iterations, and 1000 voxels used to estimate hyperparameters. A quadratic first level model was used to characterize Eddy current-related spatial distortion, with *Q*-space coordinates forcefully assigned to shells. Field offset was attempted to be separated from subject movement, and shells were subsequently aligned post-eddy, with *Eddy*’s outlier replacement enabled^147^. Data were grouped by slice, only including values from slices determined to contain at least 250 intracerebral voxels. Final interpolation was performed using the Jacobian modulation (*jac*) method.

Based on the estimated susceptibility distortion, an unwarped *b*=0 reference was calculated for a more accurate co-registration with the anatomical reference. Framewise displacement (FD) was calculated as a confounding time-series based on the preprocessed DWI, using the implementation in *Nipype* (following the definitions by Ref.^148^). The head-motion estimates calculated in the correction step were also placed within the corresponding confounds file. Slicewise cross correlation was also calculated. The DWI time-series were resampled to ACPC, generating a preprocessed DWI run in ACPC space with 1.8mm isotropic voxels. Many internal operations of QSIPrep use *Nilearn* 0.10.1^149^ and *Dipy*^150^.

#### Diffusion MRI post-processing

Following preprocessing of structural and diffusion images, we performed tract reconstruction to obtain fractional anisotropy (FA) measures in WM tracts using *QSIRecon* 1.1.0^137^, which is based on *Nipype* 1.9.1^138,139^. Brain masks from *antsBrainExtraction* were used in all subsequent reconstruction steps.

Diffusion orientation distribution functions (ODFs) were first reconstructed using generalized q-sampling imaging (GQI)^151^, with a ratio of mean diffusion distance of 1.25 in *DSI Studio* (https://dsi-studio.labsolver.org, version Chen, 94b9c79)^100^. The reconstructed ODFs were then used as a basis to run automatic tractography in *DSI Studio* using the *AutoTrack* algorithm^69^. *AutoTrack* performs non-linear registration of subject data to MNI152NLin2009bAsym space, and places seeds for deterministic tractography in any voxel within an atlas-defined tract volume. It then uses pathway recognition based on the tractography atlas to filter out false and unrelated streamlines. *AutoTrack* attempted to reconstruct a total of 67 tracts; due to changes in the *DSI Studio* atlas over time, 32 overlapped with the tracts set used for population-level analyses in the present study. *AutoTrack* parameters were set as follows: distance tolerance was set to 22, 26, or 30 mm (evaluating all three and selecting one per tract), the track-to-voxel ratio to 2.0, and the yield rate to 10^-6^. *DSI Studio* was also used to generate FA maps for all participants. Diffusion tensors were fitted using default parameters, which involves thresholding b-values lower than 1750 s/mm² for calculating FA to avoid poor fitting results. We used the tabular derivatives of *QSIRecon* containing tract-wise mean scalar measures to summarize tract mean FA values for each tract, resulting in a participants-by-tracts matrix (1145×32).

#### Sample construction

In line with recent work using this dataset^23,89^, exclusion criteria were applied successively as follows: the initial sample of *N*=1,368 included participants with T1w images, field maps, and non-variant diffusion imaging scans^152^ with similar acquisition parameters. Medical health history led to the exclusion of *N*=118 participants due to medical conditions affecting brain function or gross neurological abnormalities. Imaging protocol quality assurance excluded *N*=25 participants due to low T1w image quality based on three expert manual raters^153^, and an additional *N*=10 individuals due to missing gradient directions in their DWI scan. Next, DWI quality control was performed using two criteria. First, we evaluated the neighborhood correlation, which measures the average pairwise spatial correlation between diffusion volumes that sample similar points in *q*-space^154^. Therefore, a low neighborhood correlation reflected poor DWI quality. As in prior work^23,89^, a neighborhood correlation threshold of 0.9 was used, resulting in the exclusion of *N*=57. Second, high head motion, defined as a framewise displacement above > 1 mm, excluded an additional *N*=13 participants. This resulted in a final study sample of *N*=1,145 available for statistical analyses.

### Individual-level statistical analysis

#### Generalized Additive Models (GAMs)

To model linear and non-linear associations between tract FA and age, generalized additive models (GAMs) were fitted using the *mgcv* package (version 1.8.39) in R 4.4.2^155–159^. A GAM was fitted separately for each tract, with mean FA as the dependent variable, age as a smooth term, and both sex and in-scanner motion (mean framewise displacement) as linear covariates. As in prior work^89,160^, the smooth term basis for age was modeled using thin plate regression splines. Restricted maximal likelihood was used for smoothing parameter selection, and a maximum basis complexity (*k*) of 3 was set to avoid overfitting. The resulting age spline, or smooth function, consists of a linear combination of weighted basis functions that capture the developmental trajectory for FA in a given tract.

Our measure of interest was the magnitude of the age effect, which as in past work^89,160^ was quantified using the partial R^2^, obtained by comparing the variance explained by the full model to a model that included covariates only. An analysis of variance (ANOVA) approach was used to compare the full and reduced models and evaluate the significance of the age effect. To assess whether cognition was associated with tract mean FA, we included the executive efficiency factor score as a linear variable in the model above. We similarly used an ANOVA comparing the full model to a model that only included age and covariates to obtain a cognition effect and perform significance testing. We used a signed version of the partial R^2^ to reflect the direction of the effects. For age models, the sign reflected the mean derivative of the age smooth term; for cognition models, the sign was based on the *t*-value associated with the linear cognition term. A positive sign indicated an overall increasing trend of FA with age (or cognition), whereas a negative sign reflected a decreasing trend. In each set of analyses, multiple comparisons were controlled for using the False Discovery Rate correction (FDR, *q* < 0.05).

#### Non-parametric permutation testing

Similarly to the assessment of associations between population-level tract annotations, the association between tract annotations—describing the hierarchical span and cognitive diversity of tracts—and tract age or cognition effects was evaluated using non-parametric permutation testing with 10,000 iterations (see “Significance Testing” section above). Correlations were FDR-corrected for multiple comparisons.

## Resource availability

### Data availability

The tract-to-region connectome is publicly available at https://brain.labsolver.org. The *Neurosynth* database is available at https://neurosynth.org. The *Cognitive Atlas* is available at https://www.cognitiveatlas.org. Cortical surface maps are available through *Neuromaps*: https://netneurolab.github.io/neuromaps.

This paper used publicly available data from the Philadelphia Neurodevelopmental Cohort, accessible from the Database of Genotypes and Phenotypes (phs000607.v3.p2) at https://www.ncbi.nlm.nih.gov/projects/gap/cgi-bin/study.cgi?study_id=phs000607.v3.p2.

### Code availability

Code used to perform the analyses is available at https://github.com/PennLINC/tractmaps.

## Acknowledgements

This study was supported by grants from the National Institute of Health: R01MH120482 (T.D.S.), R01MH113550 (T.D.S.), R01MH112847 (R.T.S. and T.D.S.), R01EB022573 (T.D.S.), RF1MH116920 (T.D.S.), R37MH125829 (T.D.S.), F31MH136685 (J.B.), R01MH089983 (R.C.G., R.E.G), R01MH117014 (R.C.G., R.E.G), R01MH096891 (R.C.G., R.E.G), R01MH132934 (A.A.B.), R01MH133843 (A.A.B.), R01MH123550 (R.T.S.), R01NS060910 (R.T.S.), R01MH123563 (R.T.S.), R01MH120174 (D.R.R.), T32MH016804 and T32MH018951 (V.J.S.). Additional support was provided by NIH U24NS130411, the AE Foundation, the Center for Artificial Intelligence and Data Science for Integrated Diagnostics at Penn, and the Penn/CHOP Lifespan Brain Institute. GS was supported by a postdoctoral fellowship from the Canadian Institutes of Health Research (CIHR).

## Author information

### Materials & Correspondence

Further information and requests regarding resources should be directed to and will be fulfilled by the lead contact, Theodore D. Satterthwaite.

### Author contributions

Conceptualization, J.B.; software: J.B., A.C.L., M.K.P., M.E.G., V.J.S., R.A.P., M.C., T.D.S.; data curation: J.B; T.T.T., M.C.; resources: J.B., R.E.G., R.C.G., T.M.M., D.R.R., F.C.Y., T.D.S.; formal analysis: J.B., G.S.; supervision: T.D.S.; validation: G.S.; investigation: J.B.; visualization: J.B.; methodology: J.B., G.S., T.S., A.A.B., R.T.S., T.D.S.; writing – original draft: J.B.; writing – review & editing: G.S., A.C.L., M.K.P., T.S., A.A.B., D.S.B., M.E.G., R.E.G., R.C.G., A.P.M., B.M.,T.M.M., D.R.R., R.T.S., V.J.S., T.T.T., F.C.Y., R.A.P., M.C., T.D.S.

### Competing interests

A.A.B. hold equity in Centile Bioscience. R.T.S. has received consulting income from Octave Bioscience and compensation for scientific reviewing from the American Medical Association.

## Supplemental information

Supplementary material contains Table S1, S2 and Figures S1-S3.

