## Supplementary Information for "Anatomical White Matter Tracts Span the Cortical Hierarchy to Support Cognitive Diversity"

Supplementary Material

### Supplementary Table 1. White matter tracts included in the study.

This table illustrates the template WM tracts included in the analyses, along with their abbreviation and tract type.

| **Tract image** | **Tract name** | **Abbreviation** | **Tract type** |
| --- | --- | --- | --- |
| 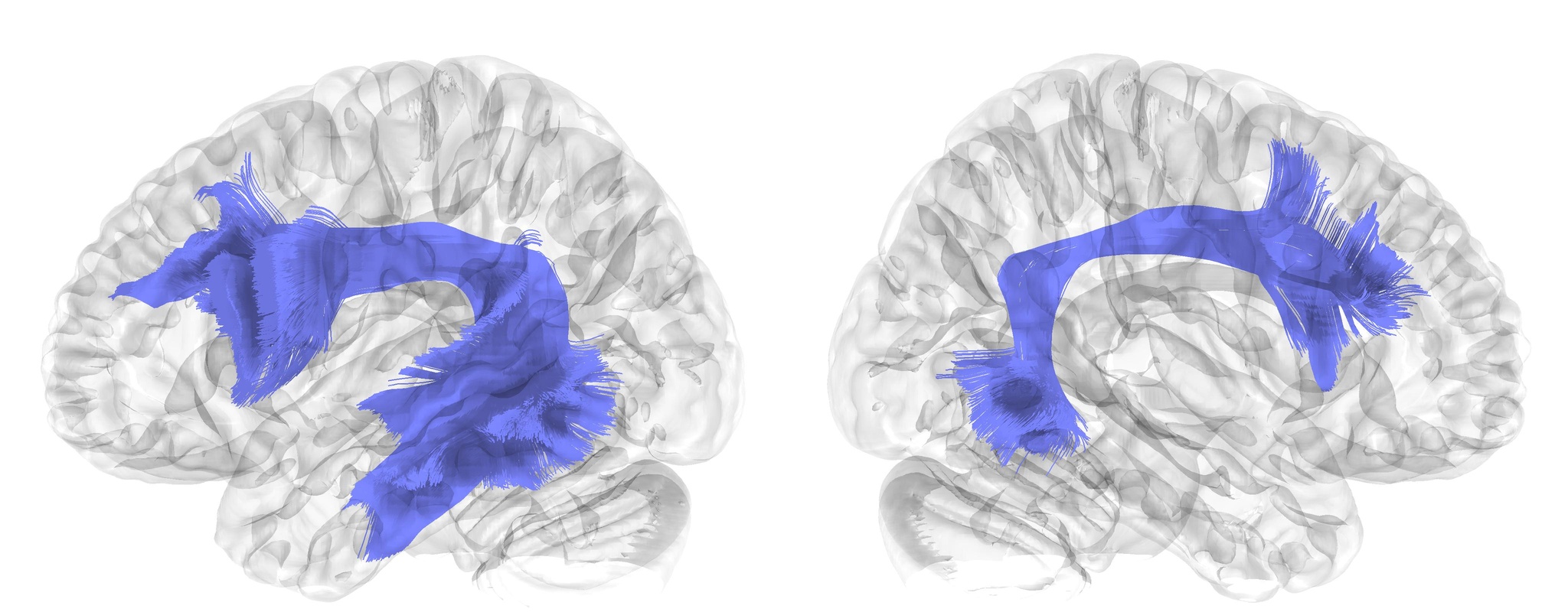 | Arcuate Fasciculus | AF | Association |
| 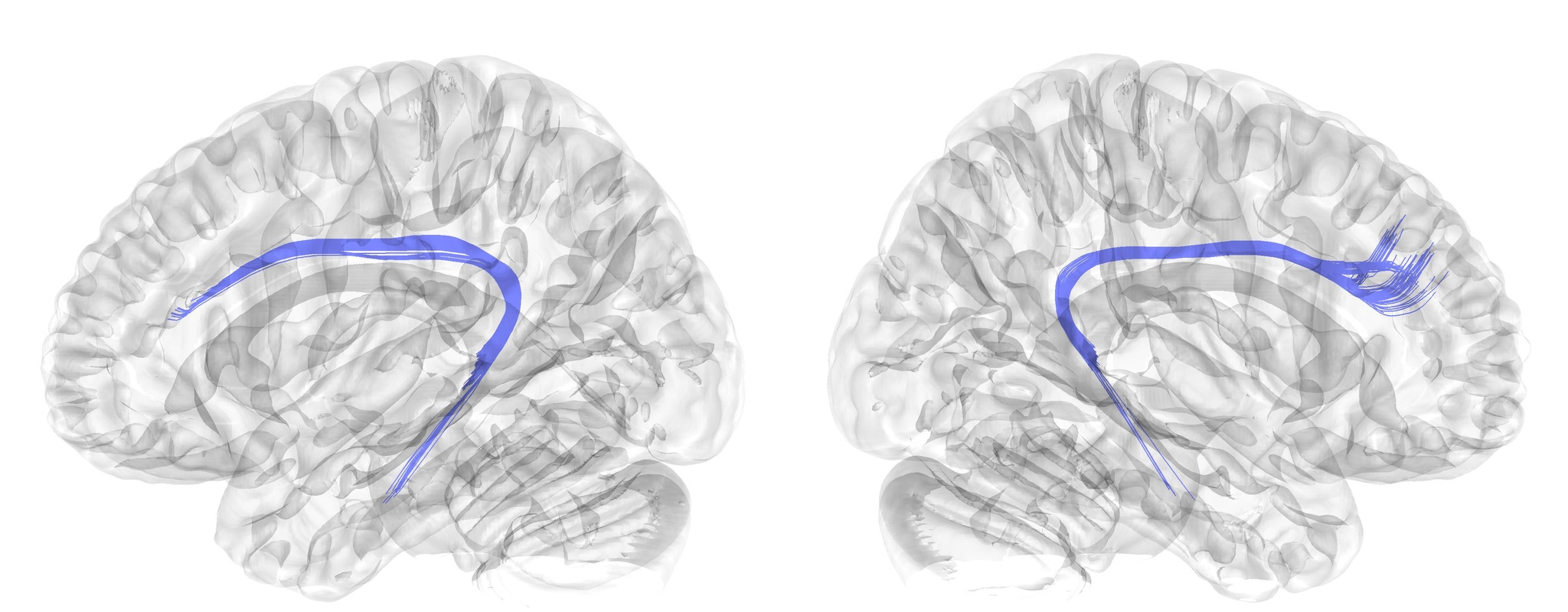 | Cingulum, Frontal Parahippocampal Segment | C_FPH | Association |
| 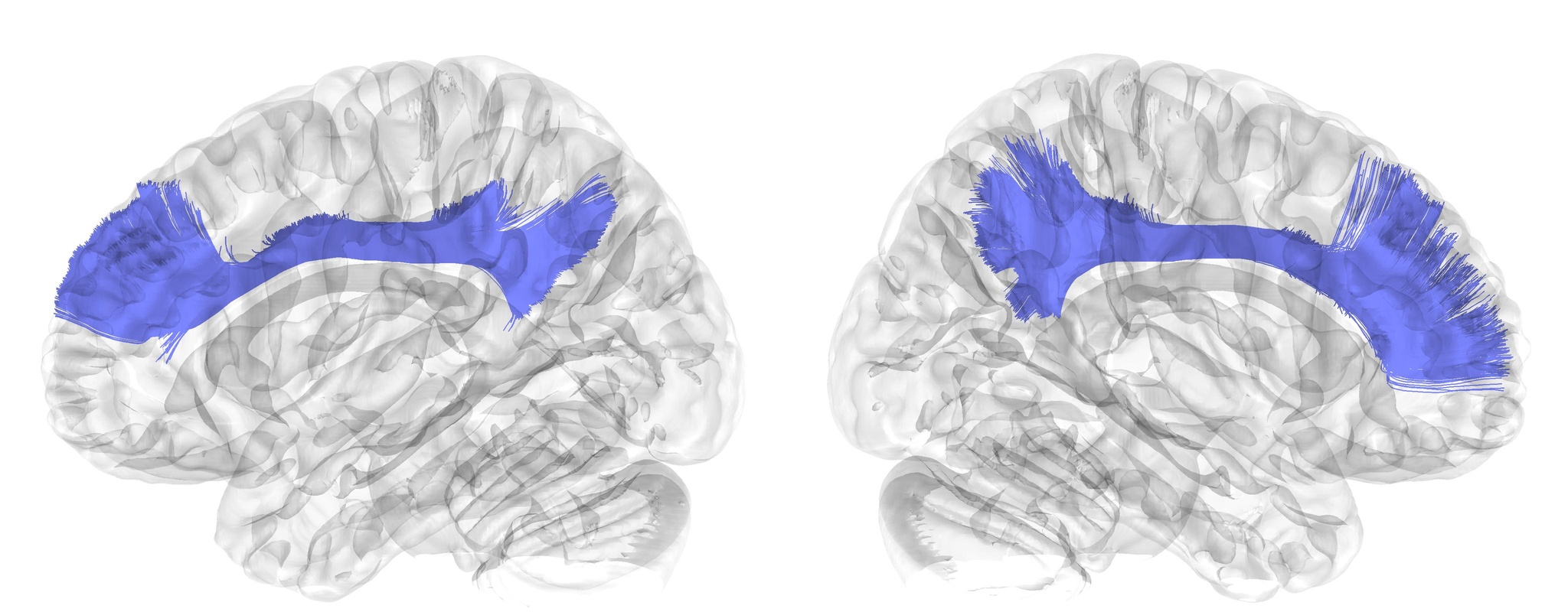 | Cingulum, Frontal Parietal Segment | C_FP | Association |
| 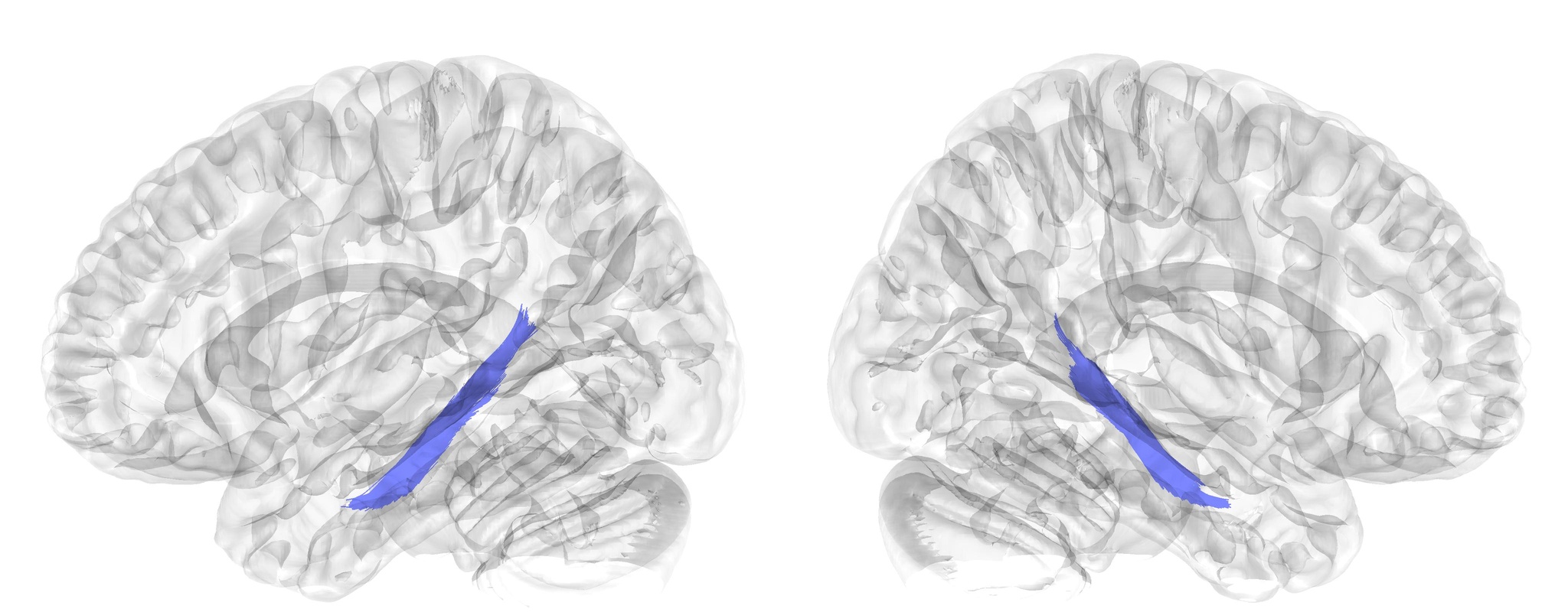 | Cingulum, Parahippocampal Segment | C_PH | Association |
| 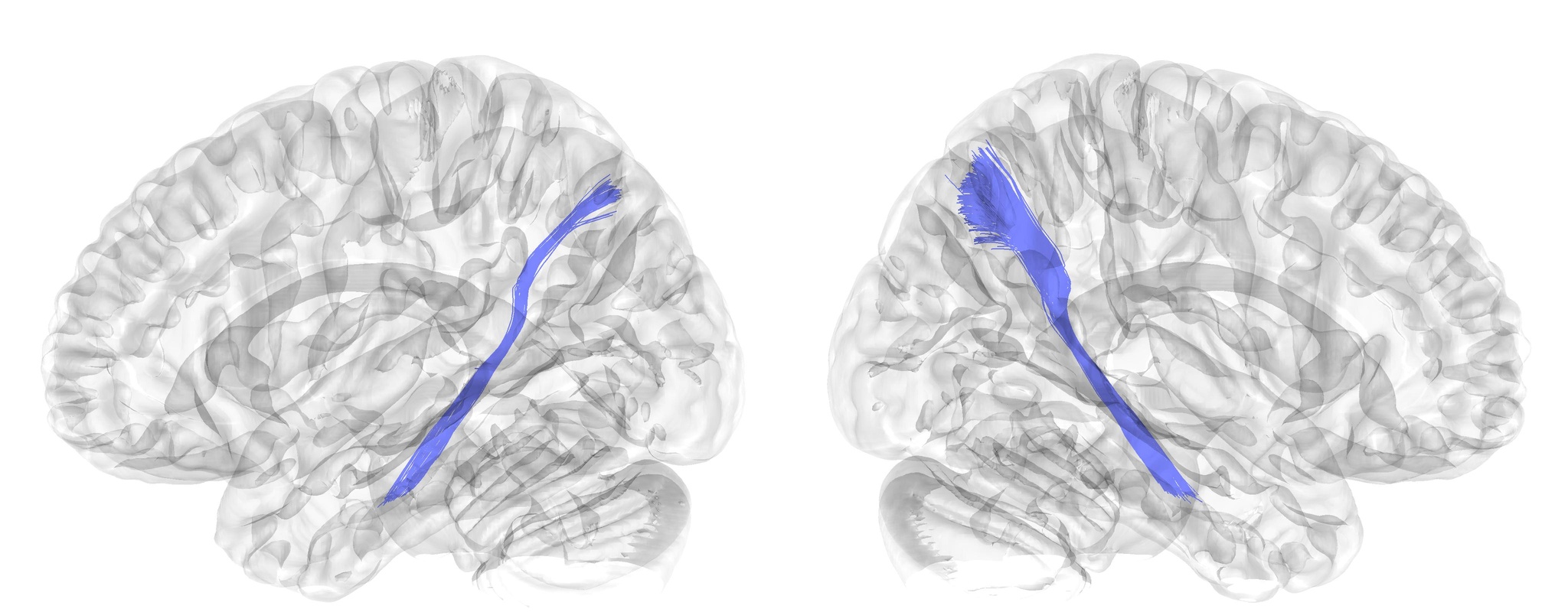 | Cingulum, Parahippocampal Parietal Segment | C_PHP | Association |
| 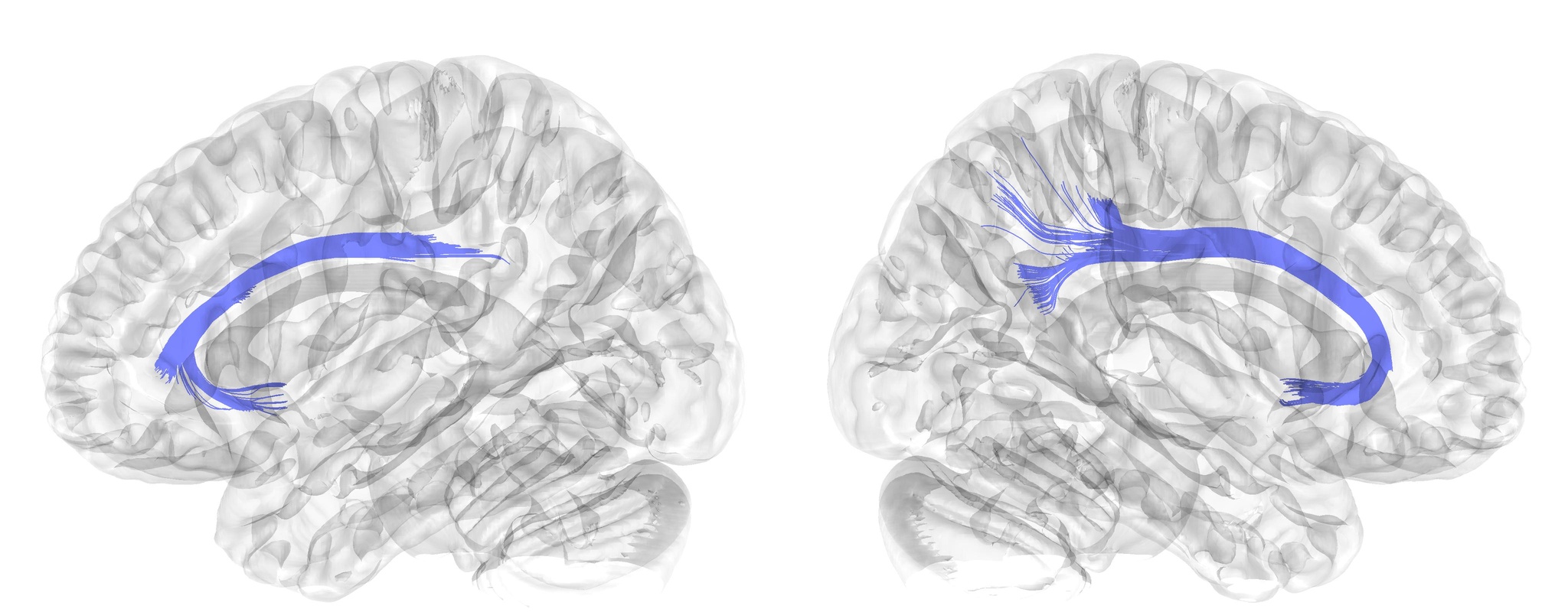 | Cingulum, Parolfactory Segment | C_R | Association |
| 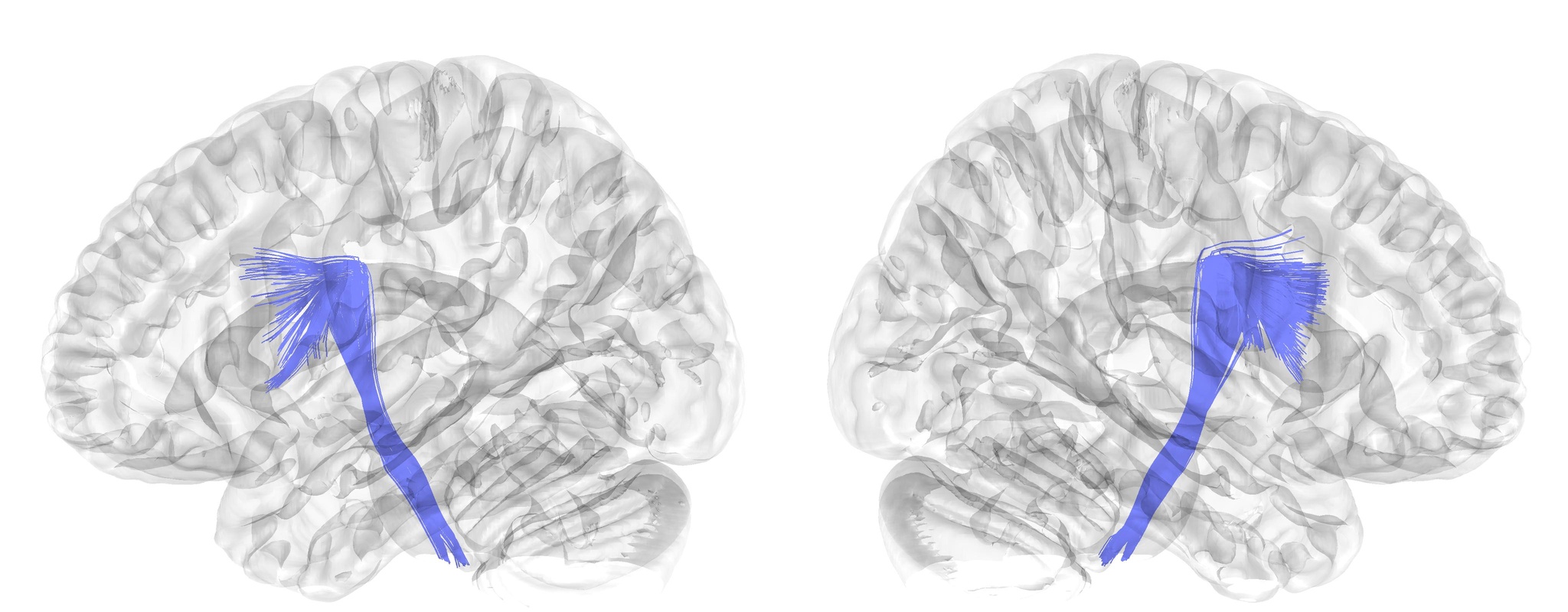 | Corticobulbar Tract | CBT | Projection |
| 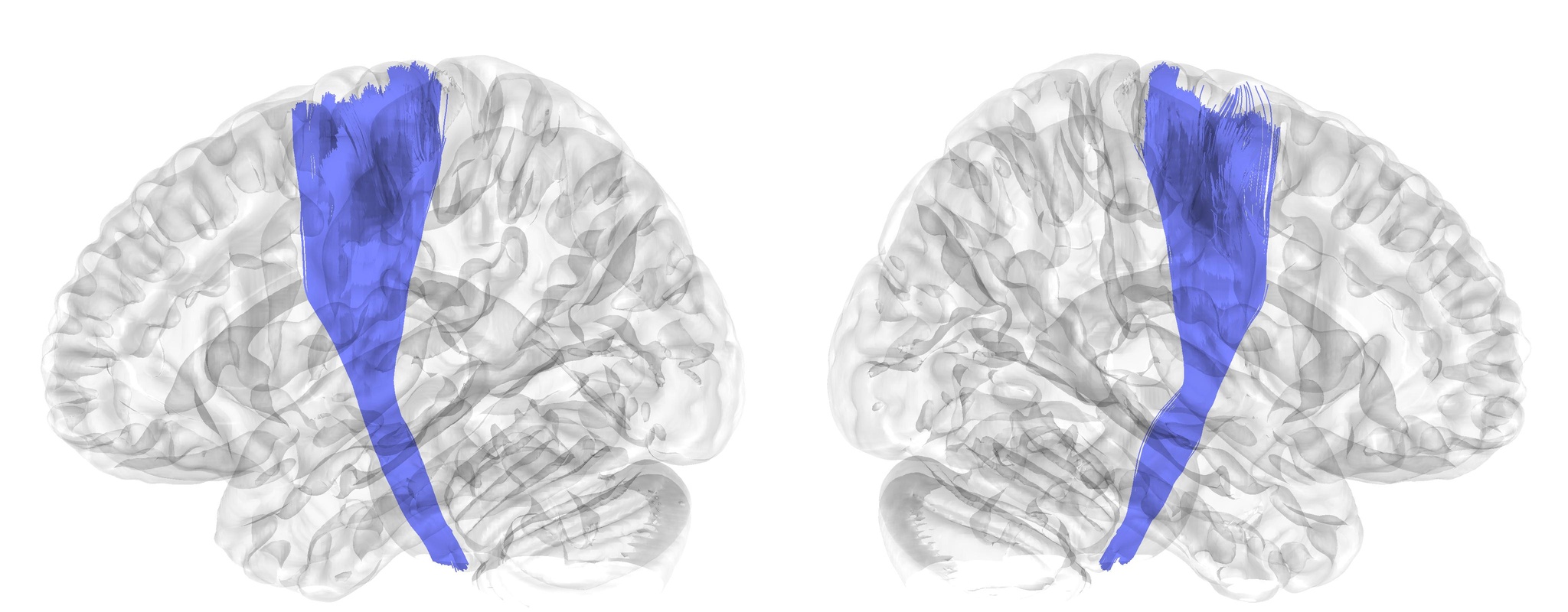 | Corticospinal Tract | CST | Projection |
| 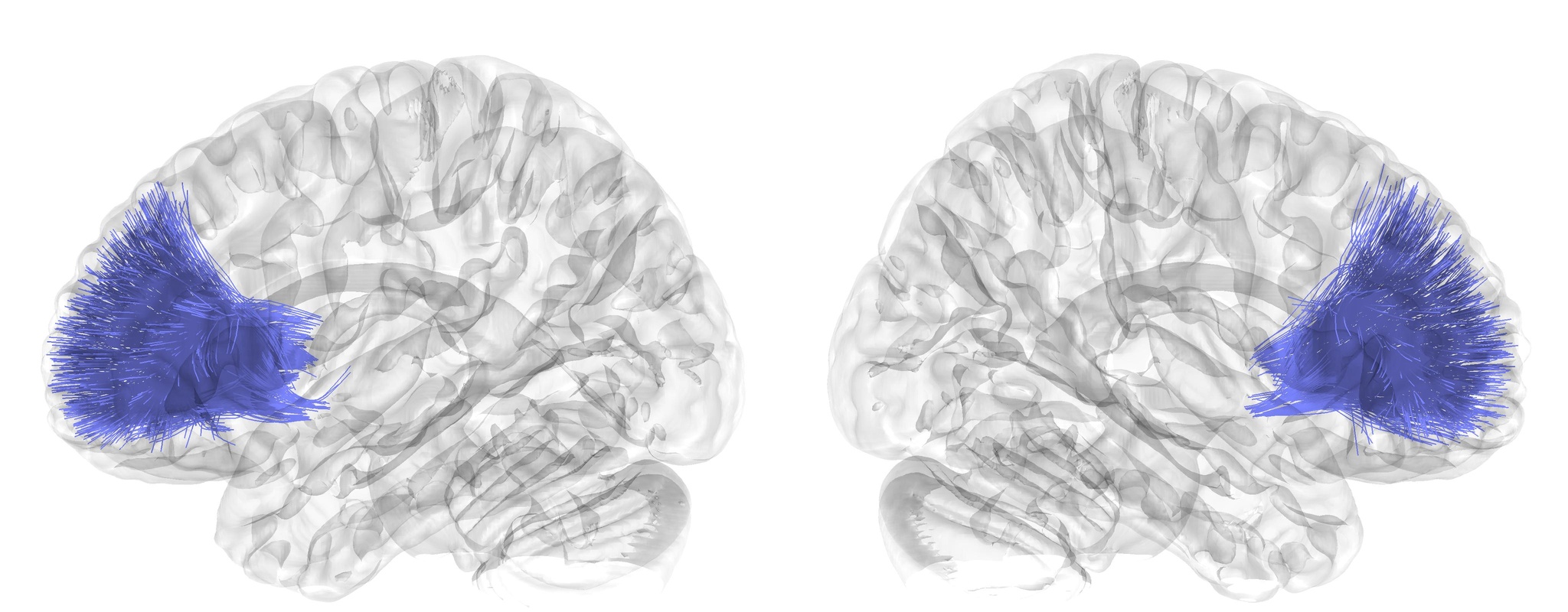 | Corticostriatal Tract, Anterior Segment | CStr_A | Projection |
| 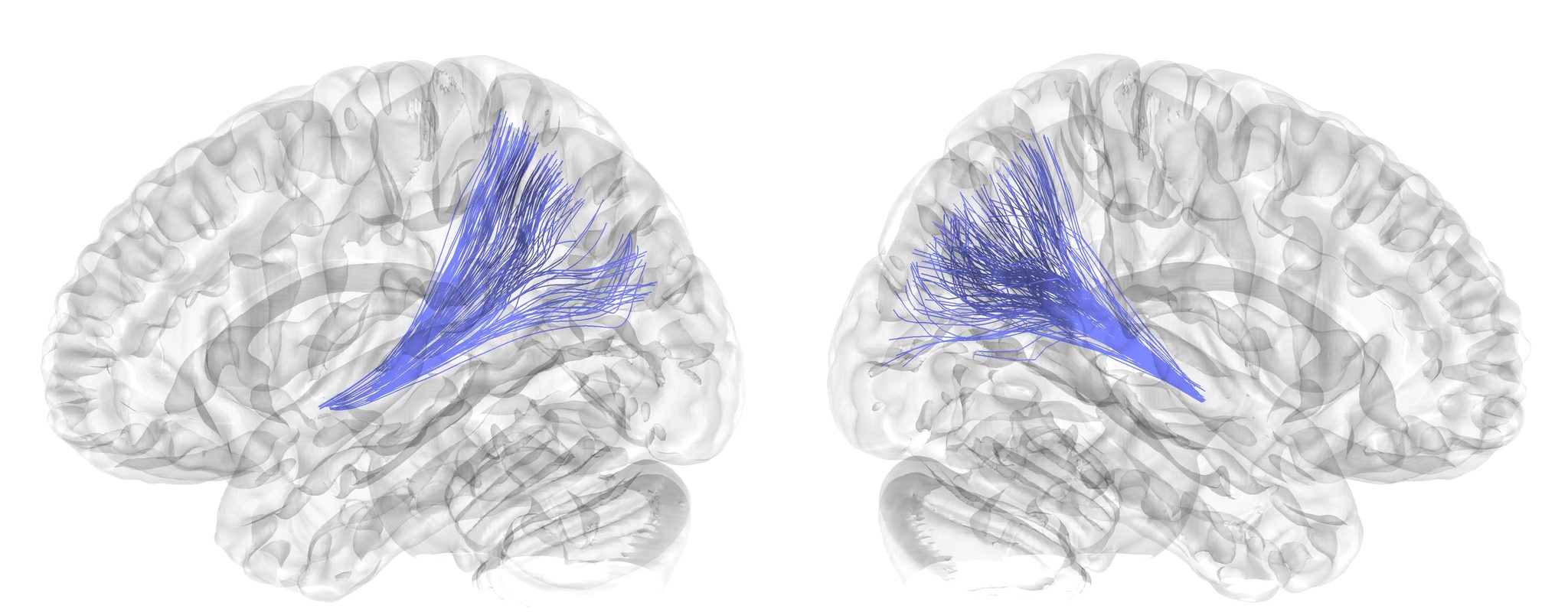 | Corticostriatal Tract, Posterior Segment | CStr_P | Projection |
| 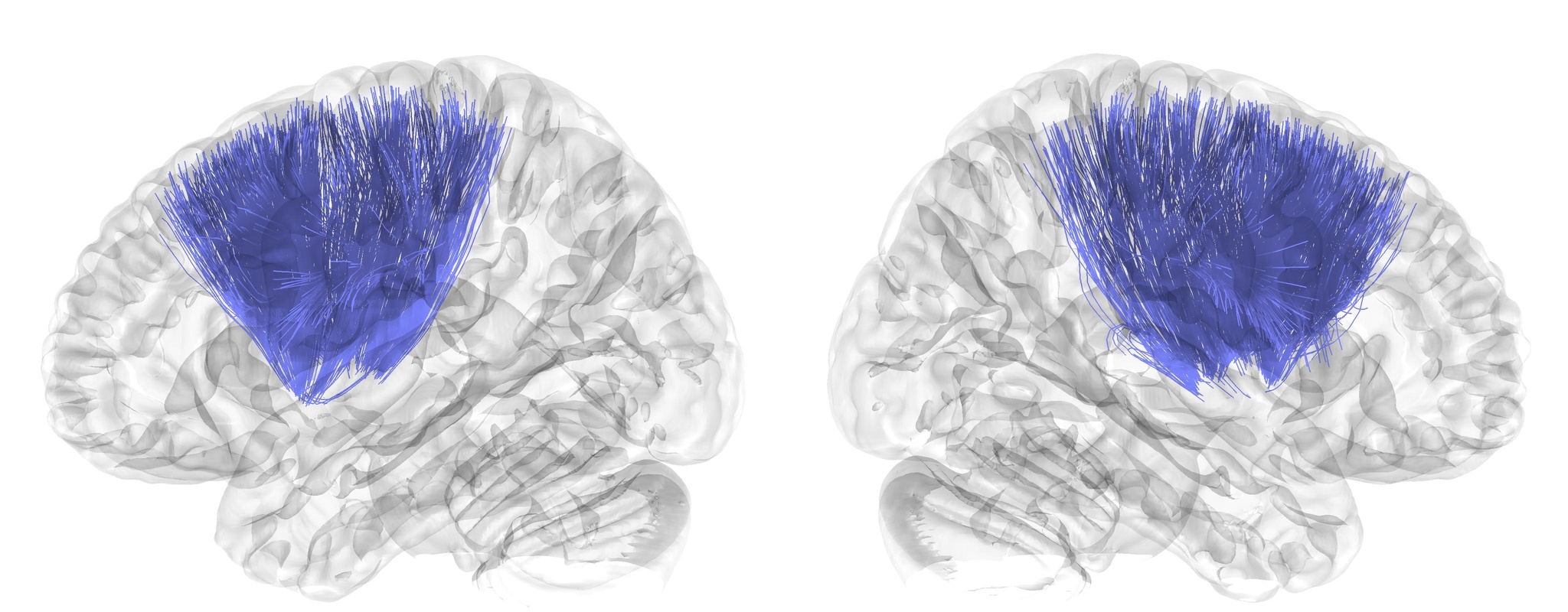 | Corticostriatal Tract, Superior Segment | CStr_S | Projection |
| 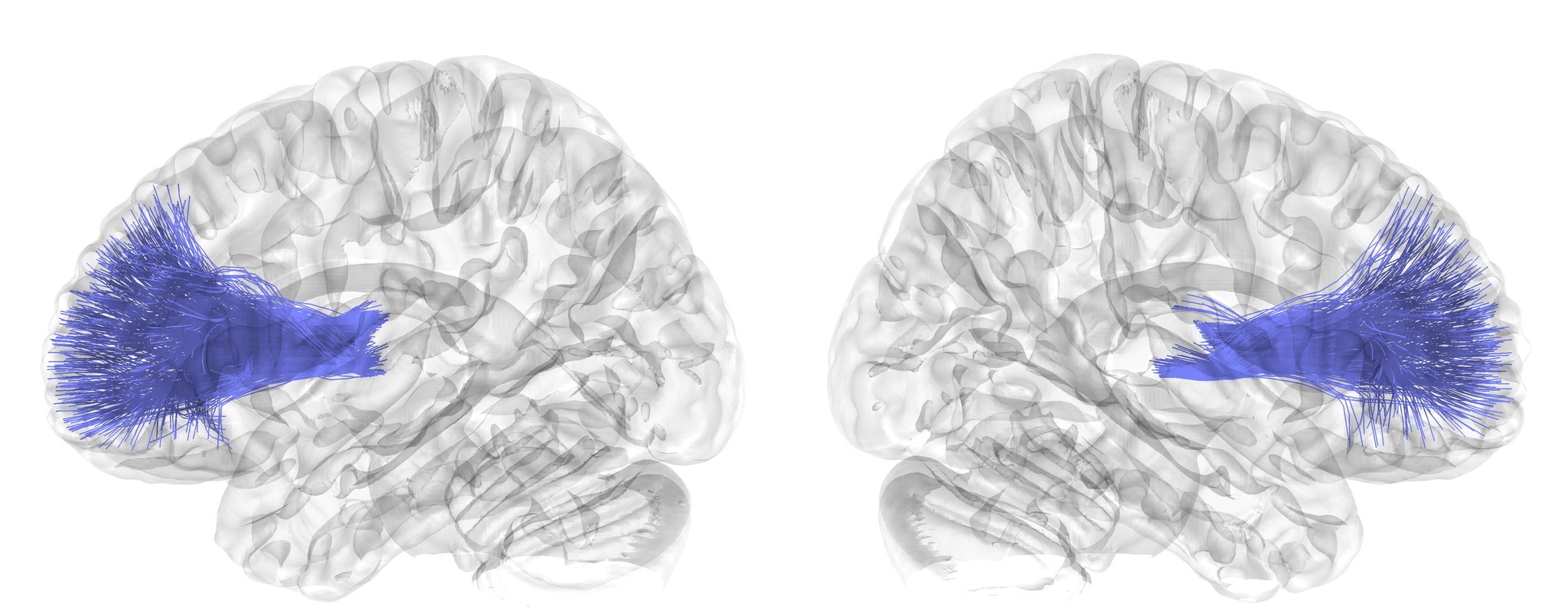 | Coricothalamic Tract, Anterior Segment | CTh_A | Projection |
| 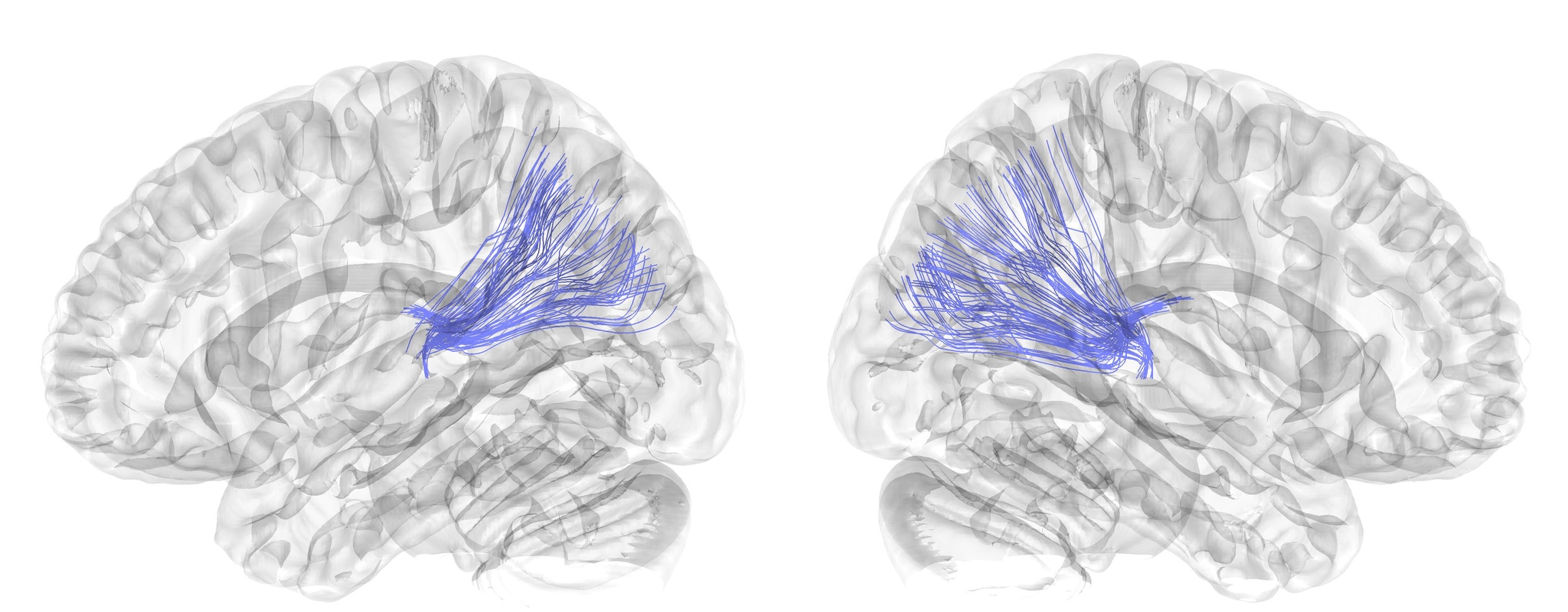 | Corticothalamic Tract, Posterior Segment | CTh_P | Projection |
| 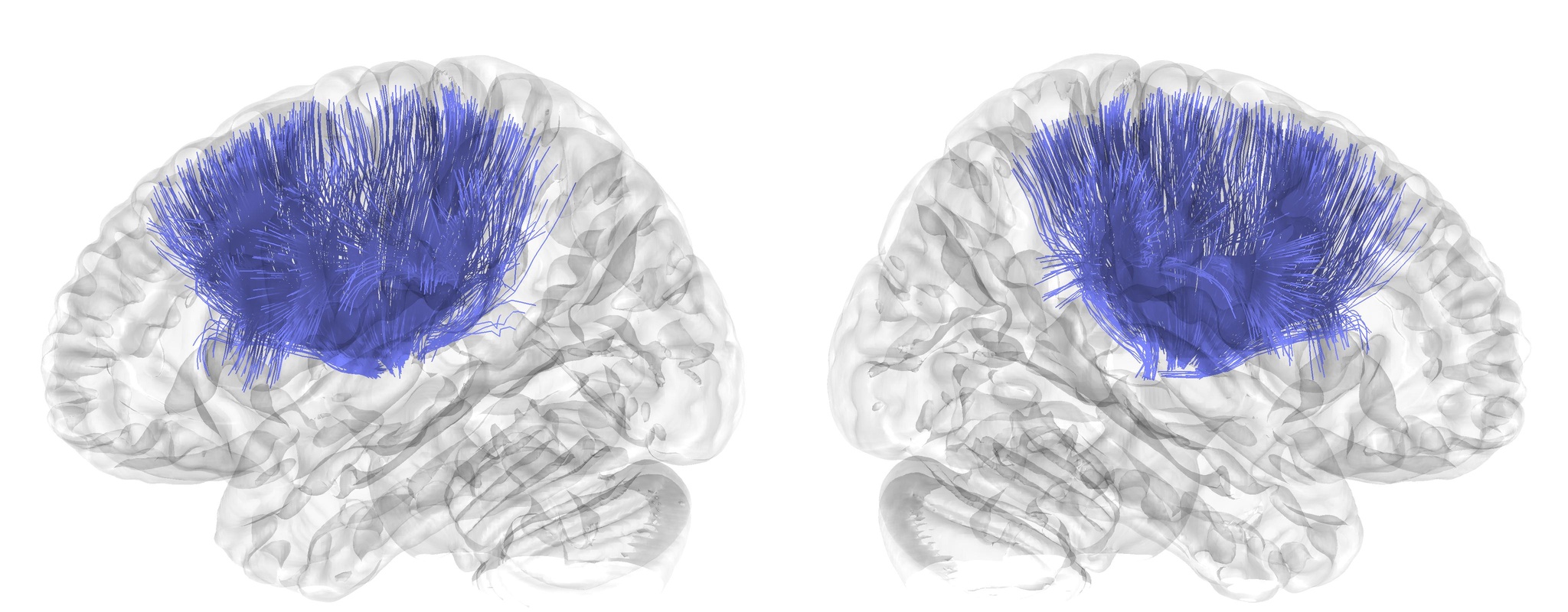 | Corticothalamic Tract, Superior Segment | CTh_S | Projection |
| 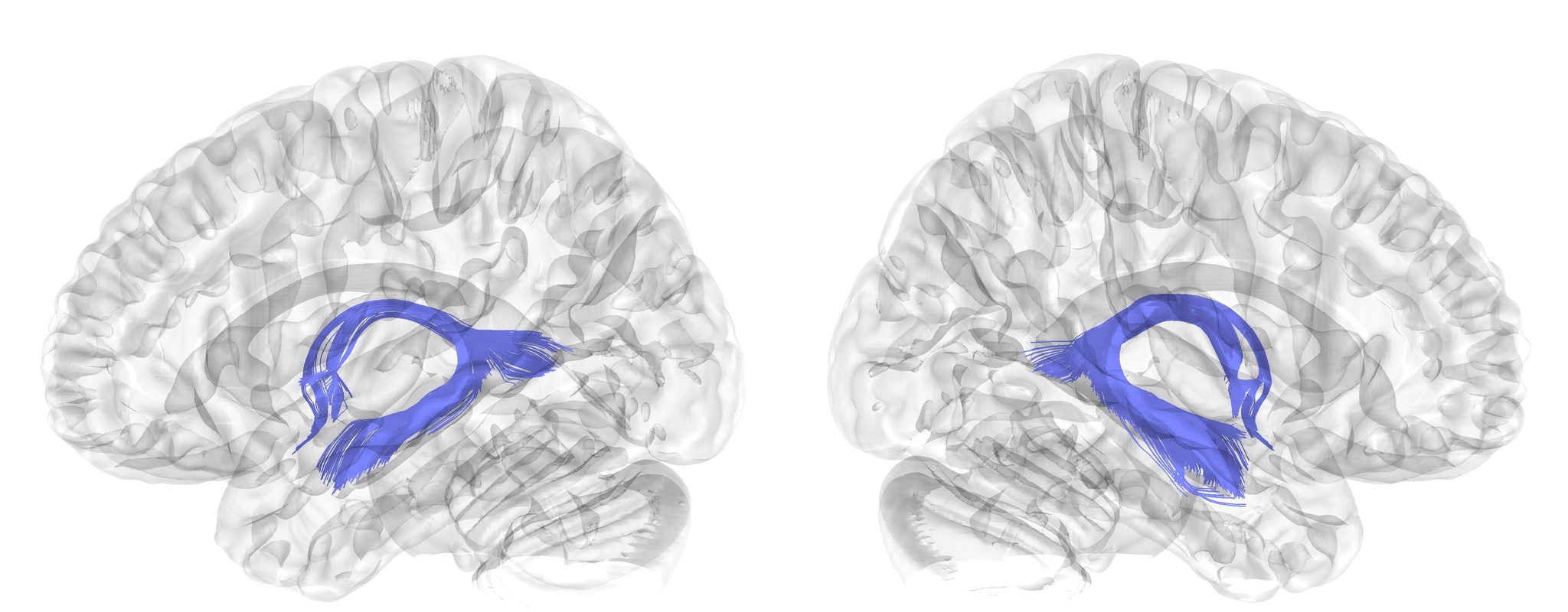 | Fornix | F | Projection |
| 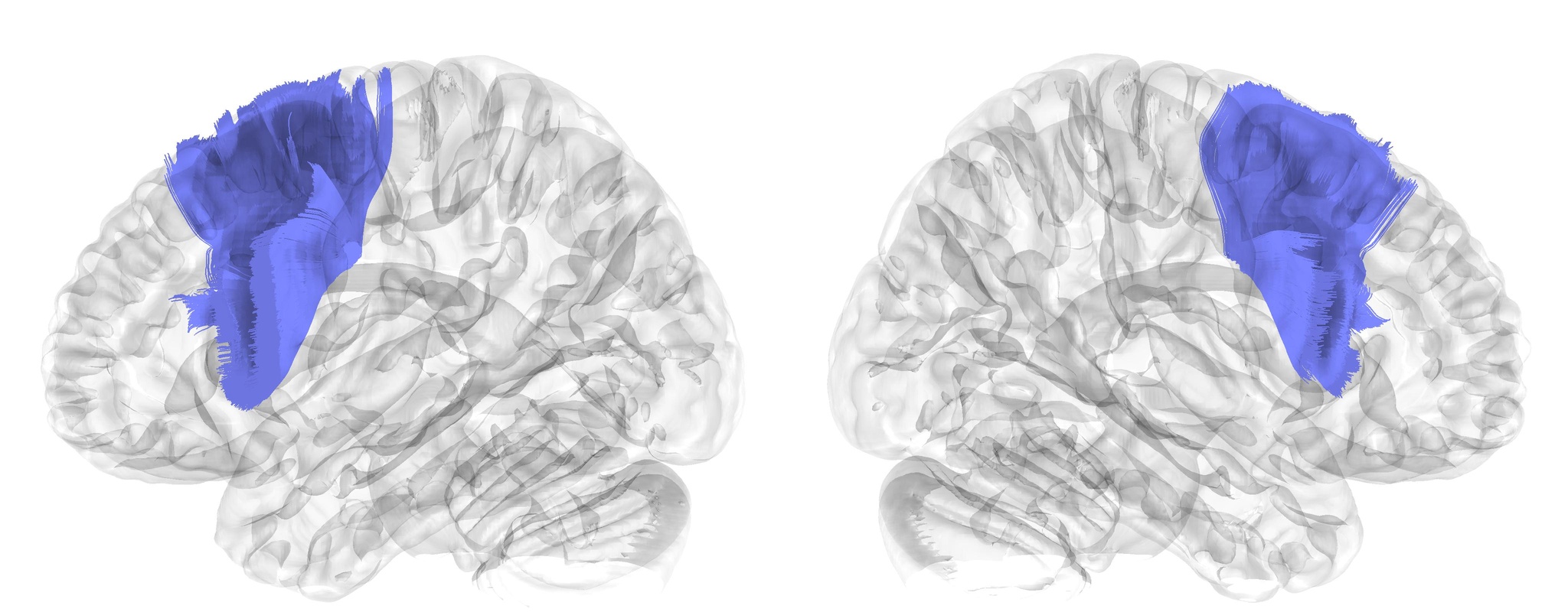 | Frontal Aslant Tract | FAT | Association |
| 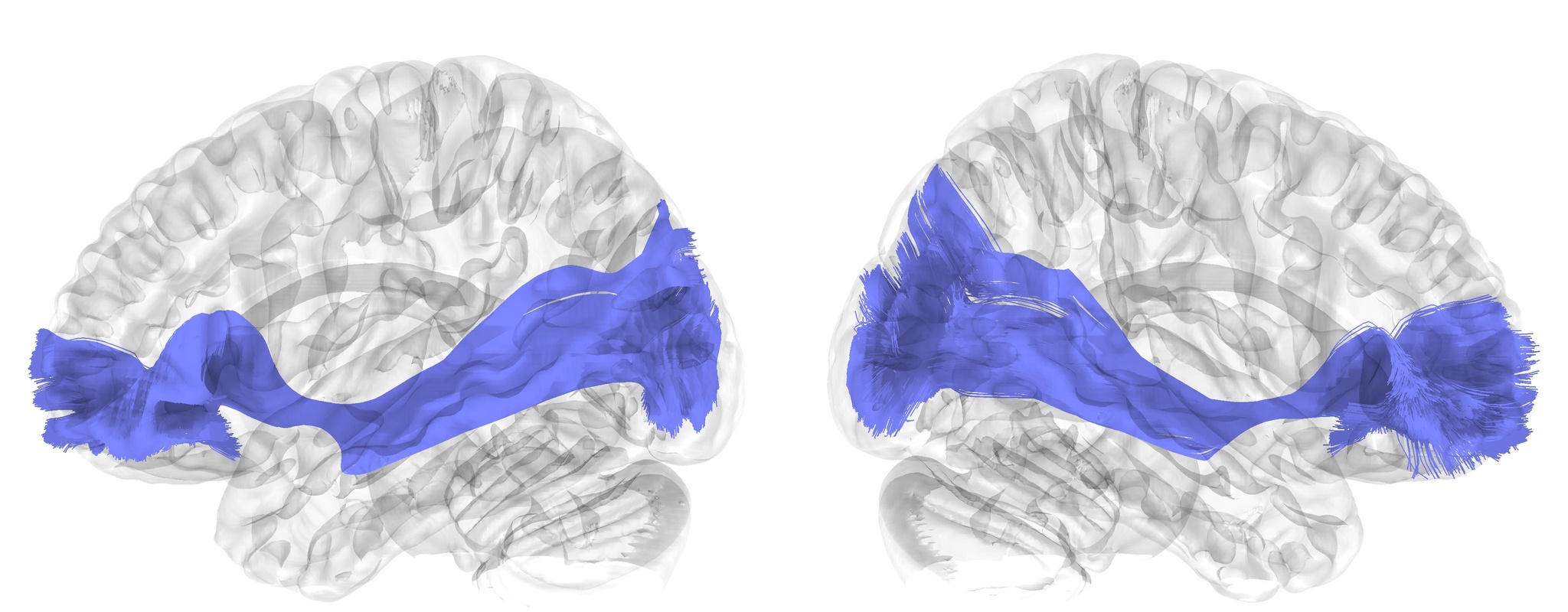 | Inferior Fronto Occipital Fasciculus | IFOF | Association |
| 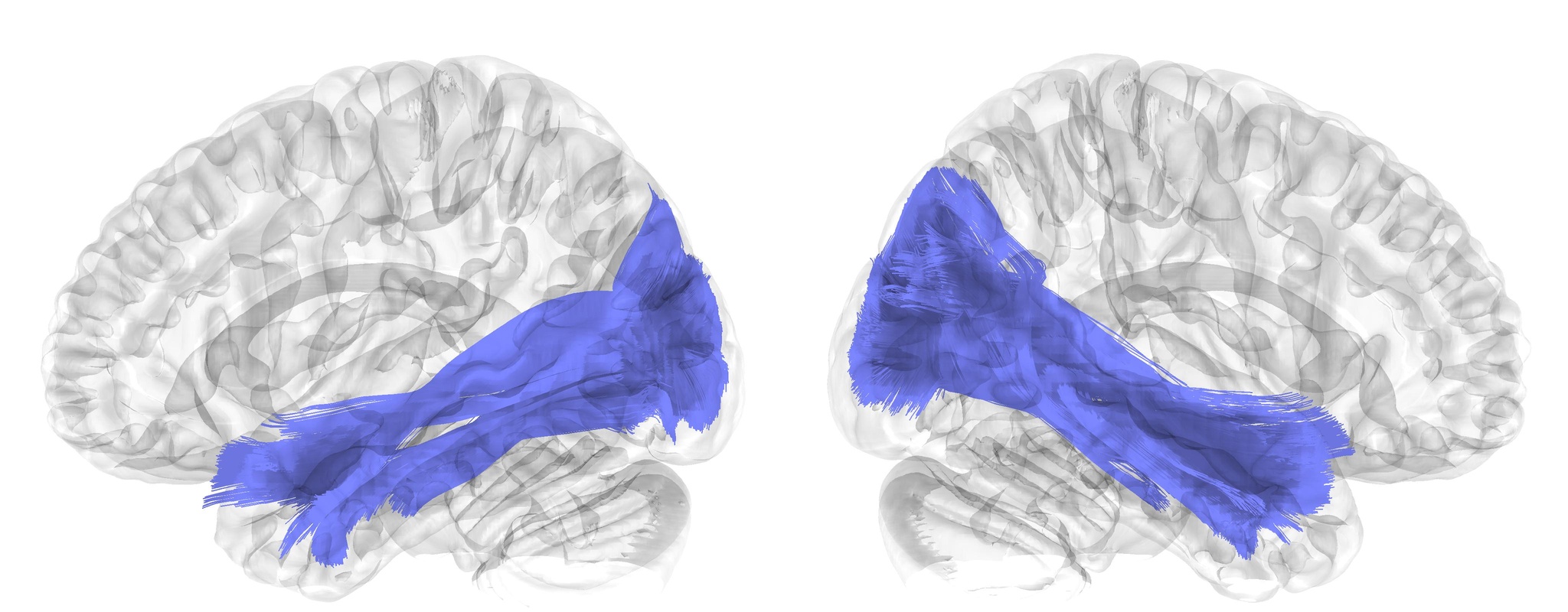 | Inferior Longitudinal Fasciculus | ILF | Association |
| 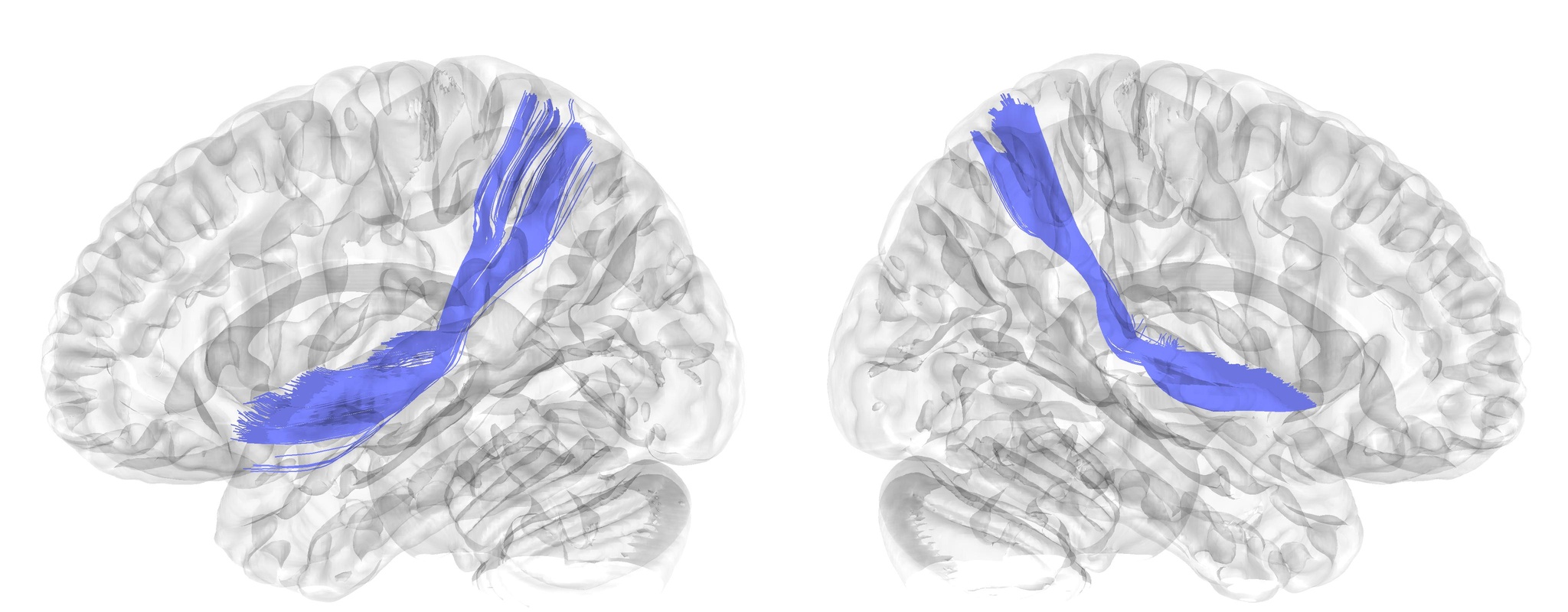 | Middle Longitudinal Fasciculus | MdLF | Association |
| 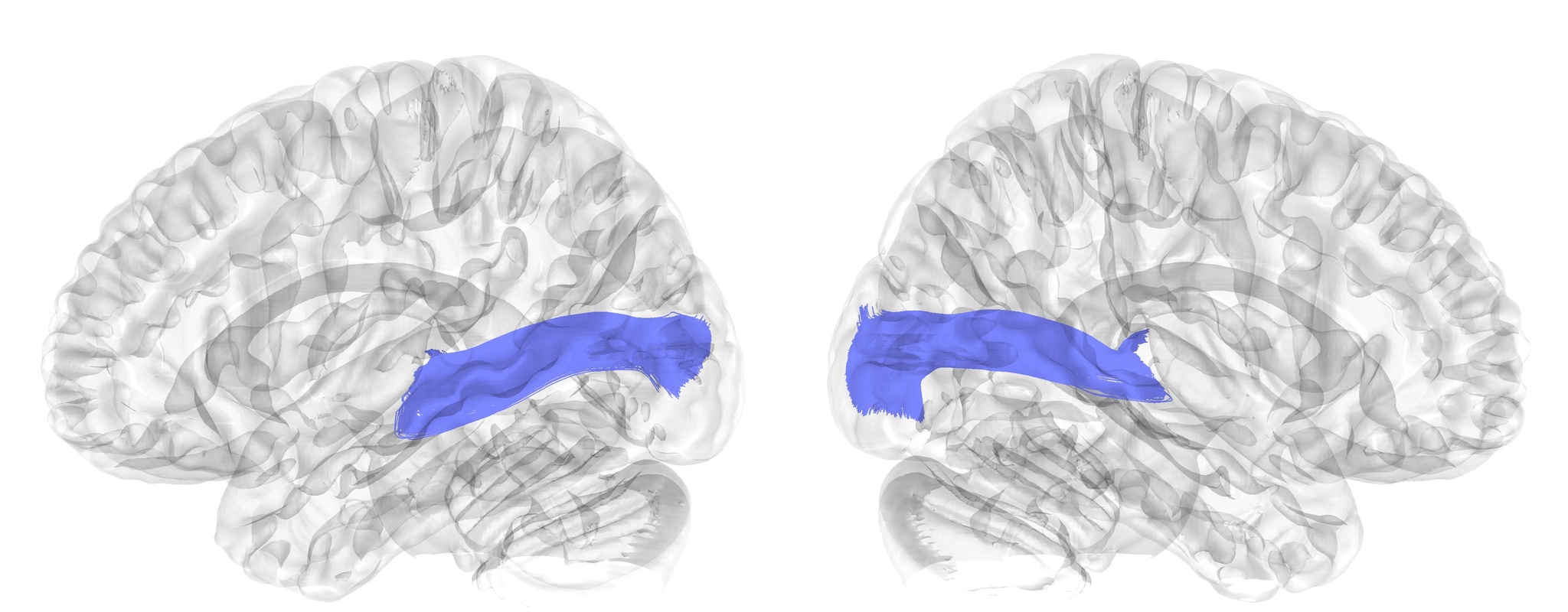 | Optic Radiation | OR | Projection |
| 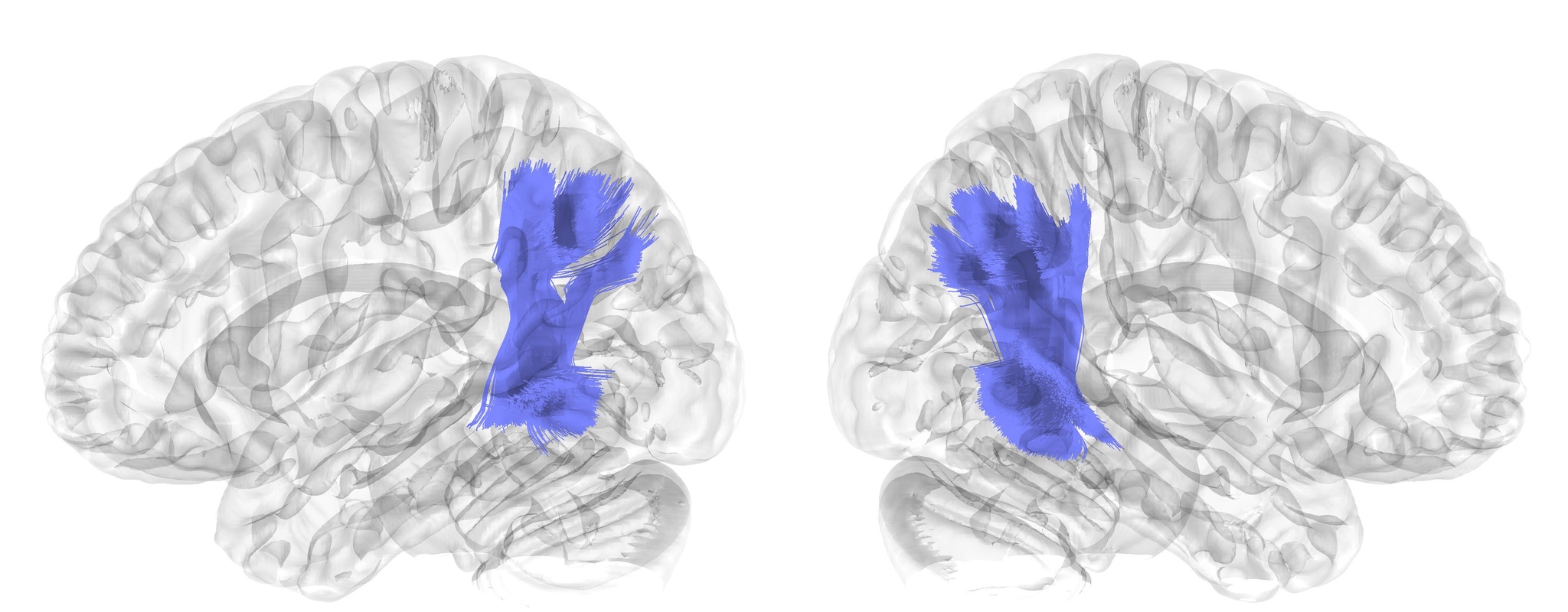 | Temporo-Parietal Aslant Tract | PTAT | Association |
| 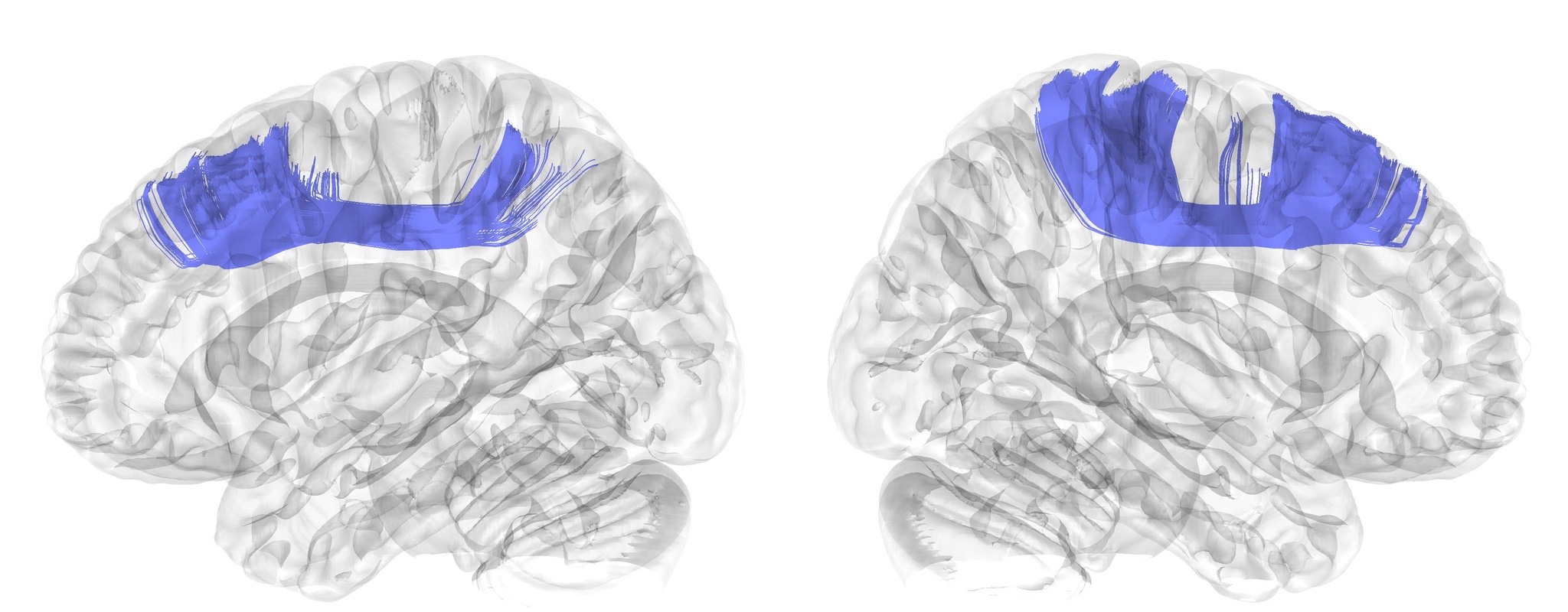 | Superior Longitudinal Fasciculus I | SLF_I | Association |
| 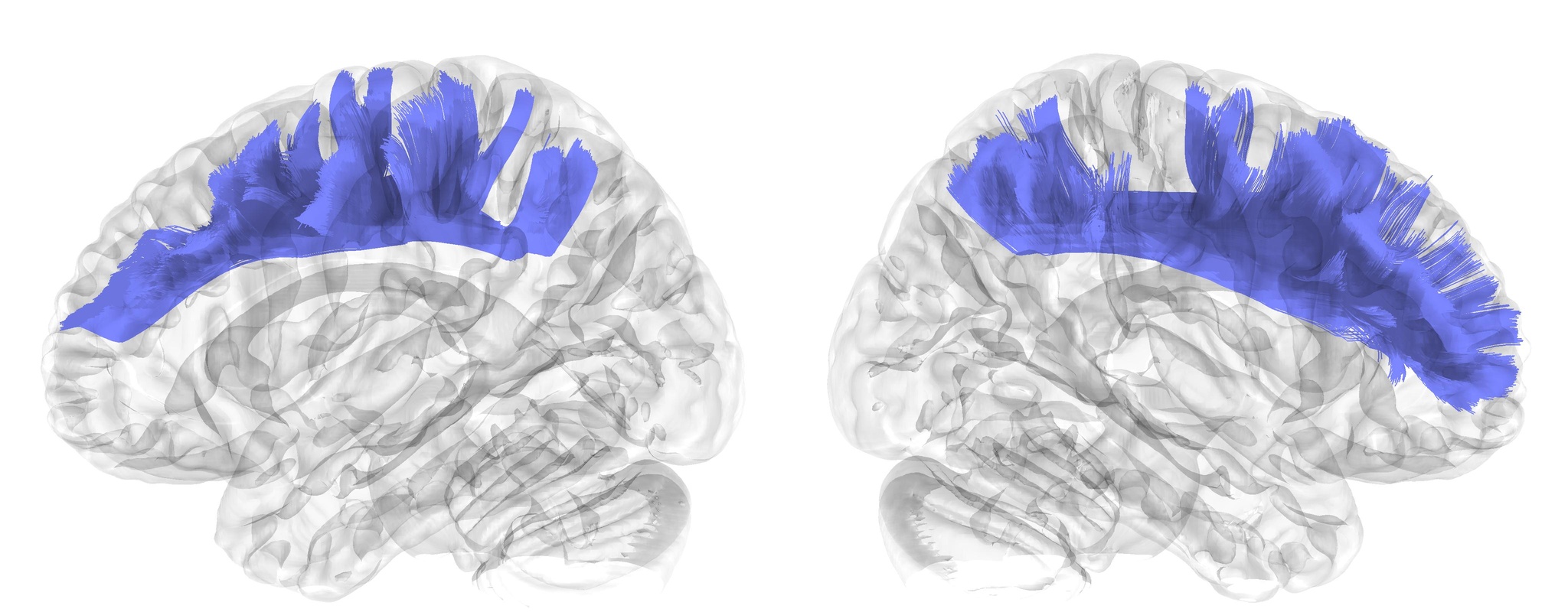 | Superior Longitudinal Fasciculus II | SLF_II | Association |
| 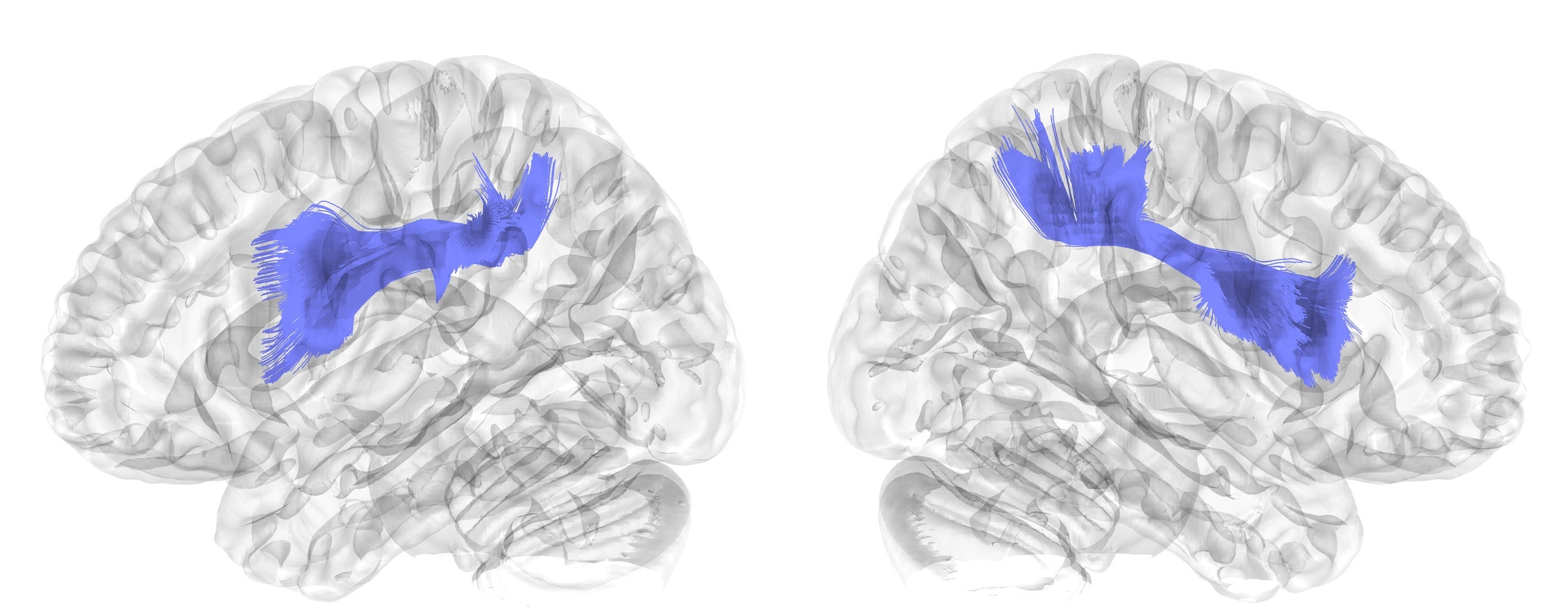 | Superior Longitudinal Fasciculus III | SLF_III | Association |
| 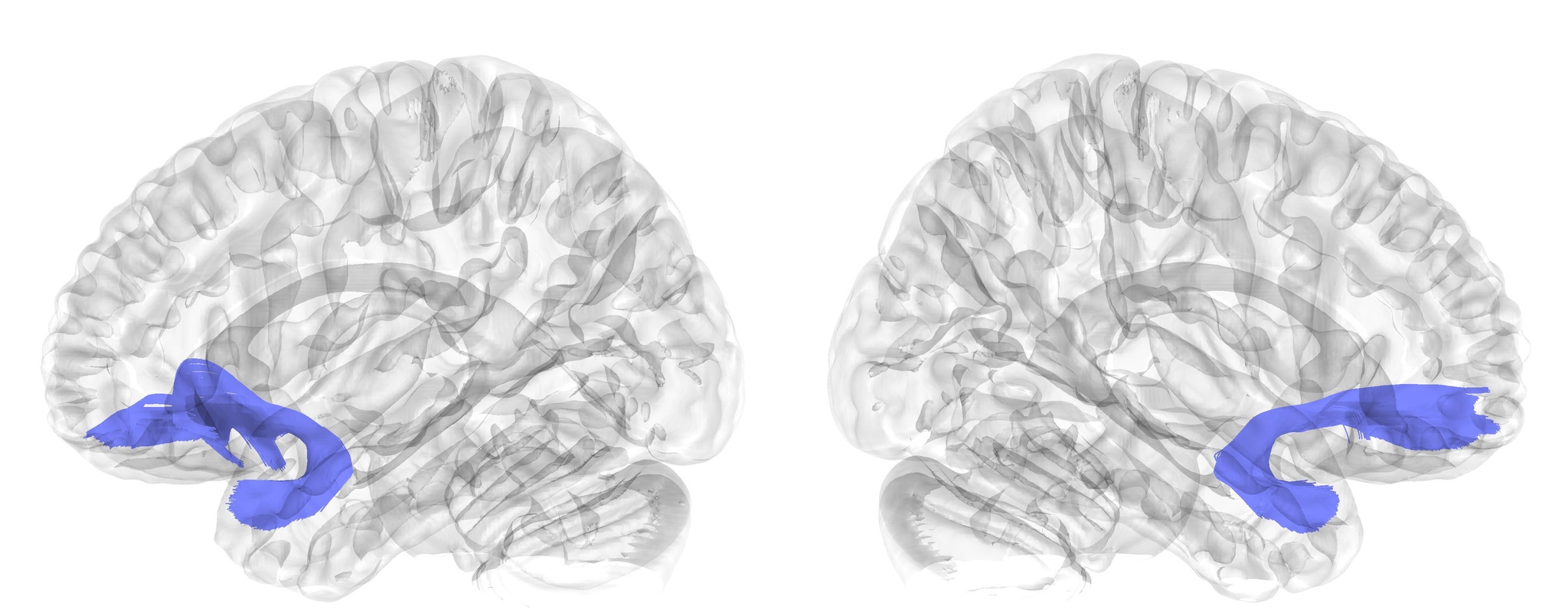 | Uncinate Fasciculus | UF | Association |
| 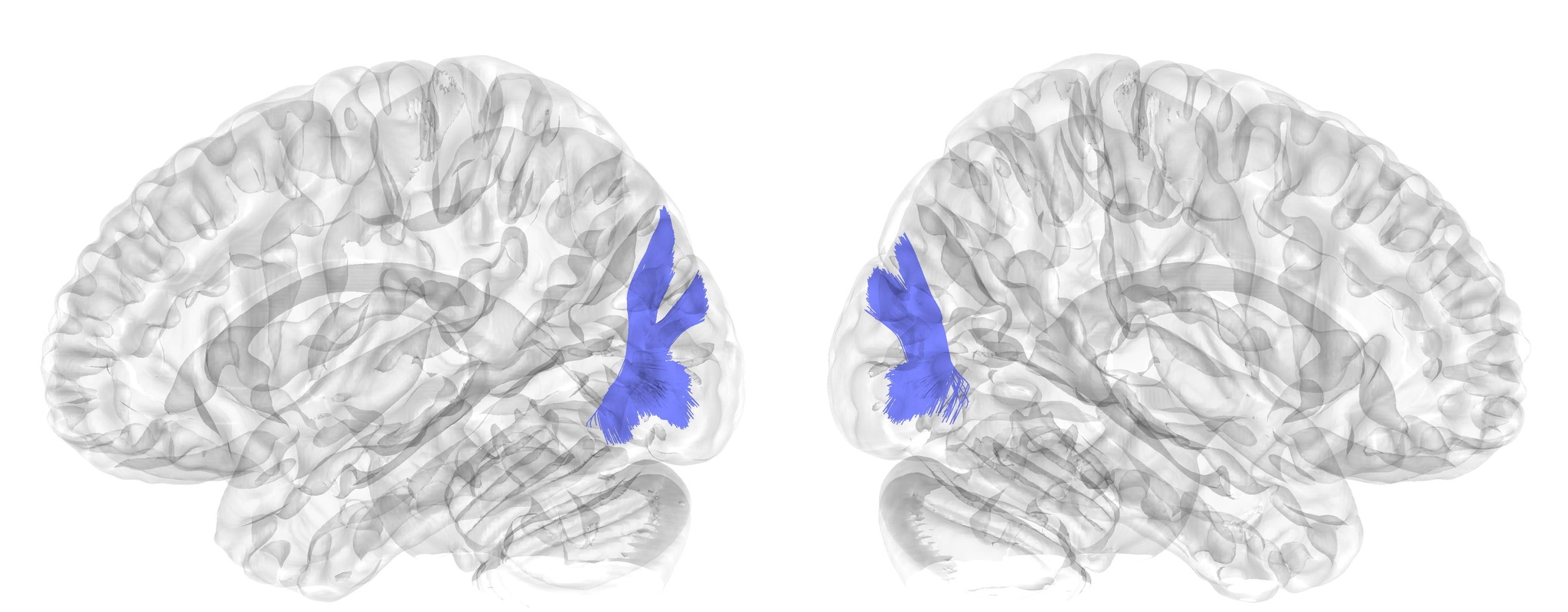 | Vertical Occipital Fasciculus | VOF | Association |

### Supplementary Table 2. Cognitive terms and categories.

This table includes cognitive terms that overlapped between the *Neurosynth*^1^ database and the *Cognitive Atlas*^2^, along with the categories they were assigned to by the *Cognitive Atlas*.

| **Cognitive term** | **Cognitive category** |
| --- | --- |
| action | action |
| motor_control | action |
| movement | action |
| response_selection | action |
| attention | attention |
| distraction | attention |
| fixation | attention |
| focus | attention |
| search | attention |
| selective_attention | attention |
| spatial_attention | attention |
| sustained_attention | attention |
| visual_attention | attention |
| categorization | decision making |
| decision | decision making |
| decision_making | decision making |
| inference | decision making |
| insight | decision making |
| intelligence | decision making |
| judgment | decision making |
| reasoning | decision making |
| risk | decision making |
| uncertainty | decision making |
| utility | decision making |
| anxiety | emotion |
| arousal | emotion |
| emotion | emotion |
| empathy | emotion |
| facial_expression | emotion |
| fear | emotion |
| mood | emotion |
| pain | emotion |
| stress | emotion |
| valence | emotion |
| cognitive_control | executive/cognitive control |
| goal | executive/cognitive control |
| inhibition | executive/cognitive control |
| maintenance | executive/cognitive control |
| manipulation | executive/cognitive control |
| monitoring | executive/cognitive control |
| planning | executive/cognitive control |
| response_inhibition | executive/cognitive control |
| updating | executive/cognitive control |
| working_memory | executive/cognitive control |
| language | language |
| language_comprehension | language |
| meaning | language |
| morphology | language |
| naming | language |
| reading | language |
| semantic_memory | language |
| sentence_comprehension | language |
| speech_perception | language |
| speech_production | language |
| verbal_fluency | language |
| word_recognition | language |
| adaptation | learning/memory |
| association | learning/memory |
| autobiographical_memory | learning/memory |
| consolidation | learning/memory |
| encoding | learning/memory |
| episodic_memory | learning/memory |
| expertise | learning/memory |
| face_recognition | learning/memory |
| familiarity | learning/memory |
| knowledge | learning/memory |
| learning | learning/memory |
| memory | learning/memory |
| memory_retrieval | learning/memory |
| priming | learning/memory |
| recall | learning/memory |
| recognition | learning/memory |
| rehearsal | learning/memory |
| reinforcement_learning | learning/memory |
| retention | learning/memory |
| retrieval | learning/memory |
| skill | learning/memory |
| expectancy | motivation |
| task_difficulty | motivation |
| addiction | other |
| anticipation | other |
| balance | other |
| belief | other |
| competition | other |
| concept | other |
| consciousness | other |
| context | other |
| coordination | other |
| eating | other |
| efficiency | other |
| effort | other |
| emotion_regulation | other |
| extinction | other |
| gaze | other |
| hyperactivity | other |
| imagery | other |
| impulsivity | other |
| induction | other |
| integration | other |
| intention | other |
| interference | other |
| listening | other |
| loss | other |
| psychosis | other |
| reward_anticipation | other |
| rule | other |
| salience | other |
| sleep | other |
| strategy | other |
| strength | other |
| timing | other |
| thought | other |
| detection | perception |
| discrimination | perception |
| localization | perception |
| mental_imagery | perception |
| multisensory | perception |
| navigation | perception |
| object_recognition | perception |
| perception | perception |
| rhythm | perception |
| transition | perception |
| visual_perception | perception |
| communication | social function |
| social_cognition | social function |

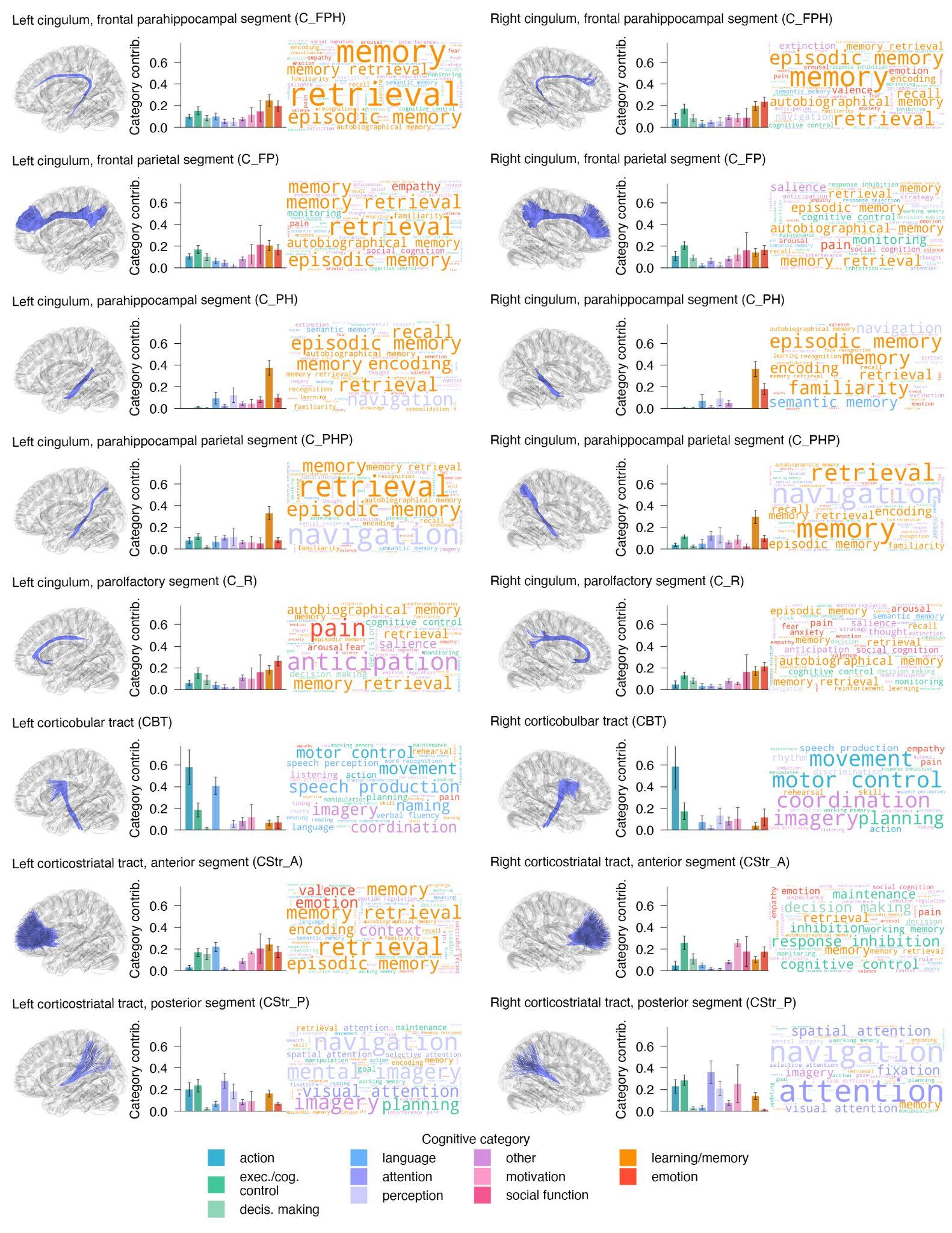

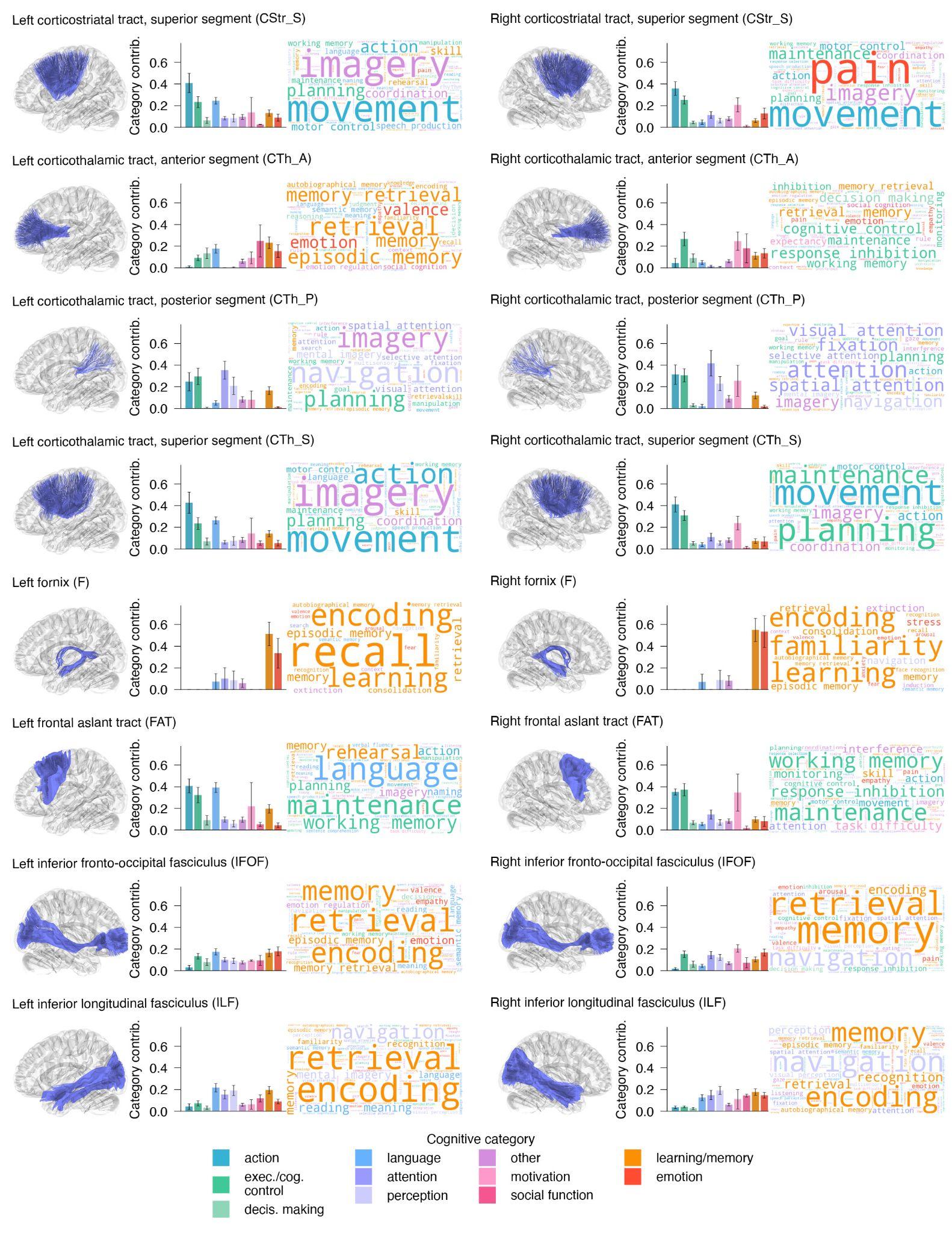

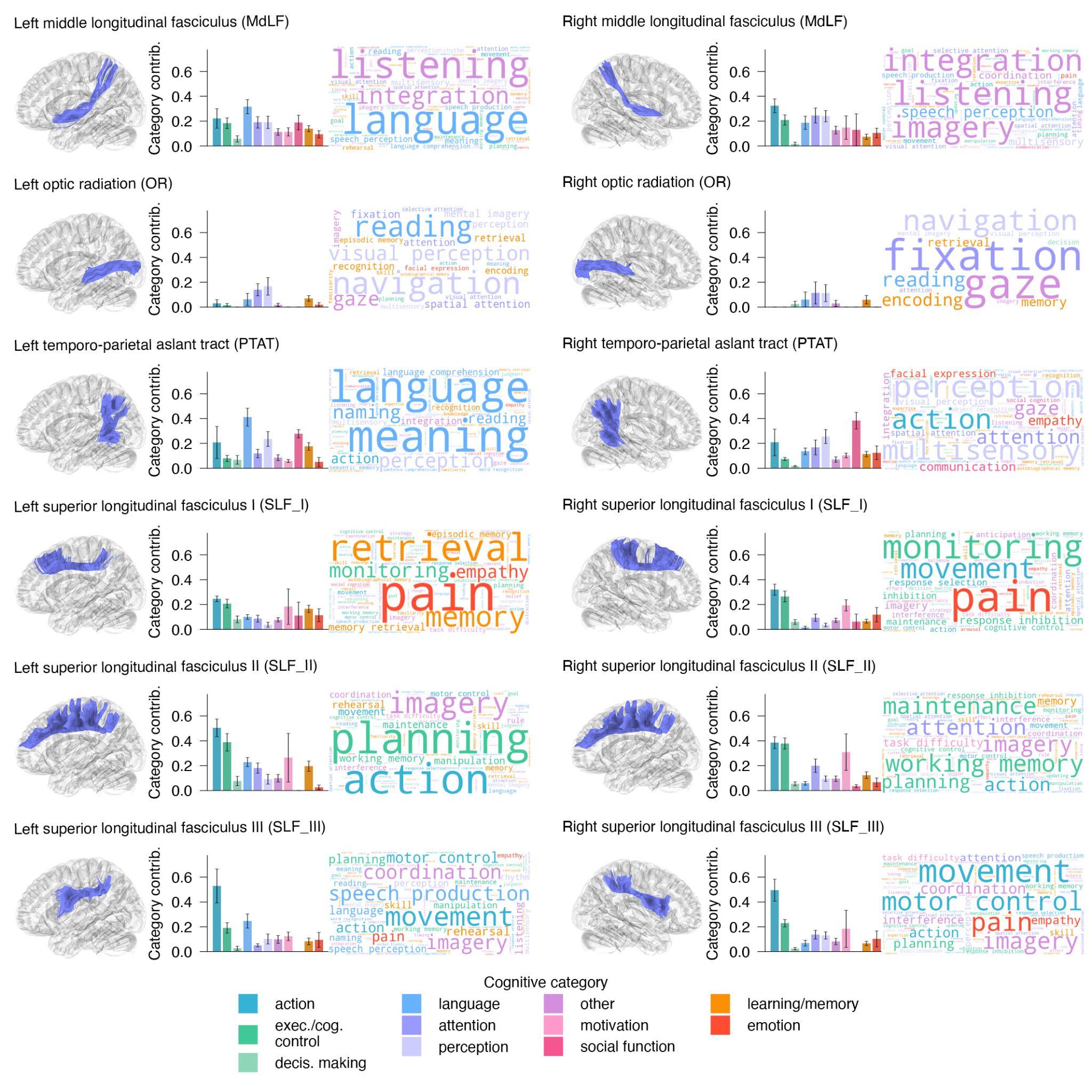

### **Supplementary Figure 1**. Cognitive profiles of all white matter tracts.

The cognitive profiles are shown for each anatomical white matter tract, with the exception of the eight exemplar tracts included in the main manuscript (Figure 4).

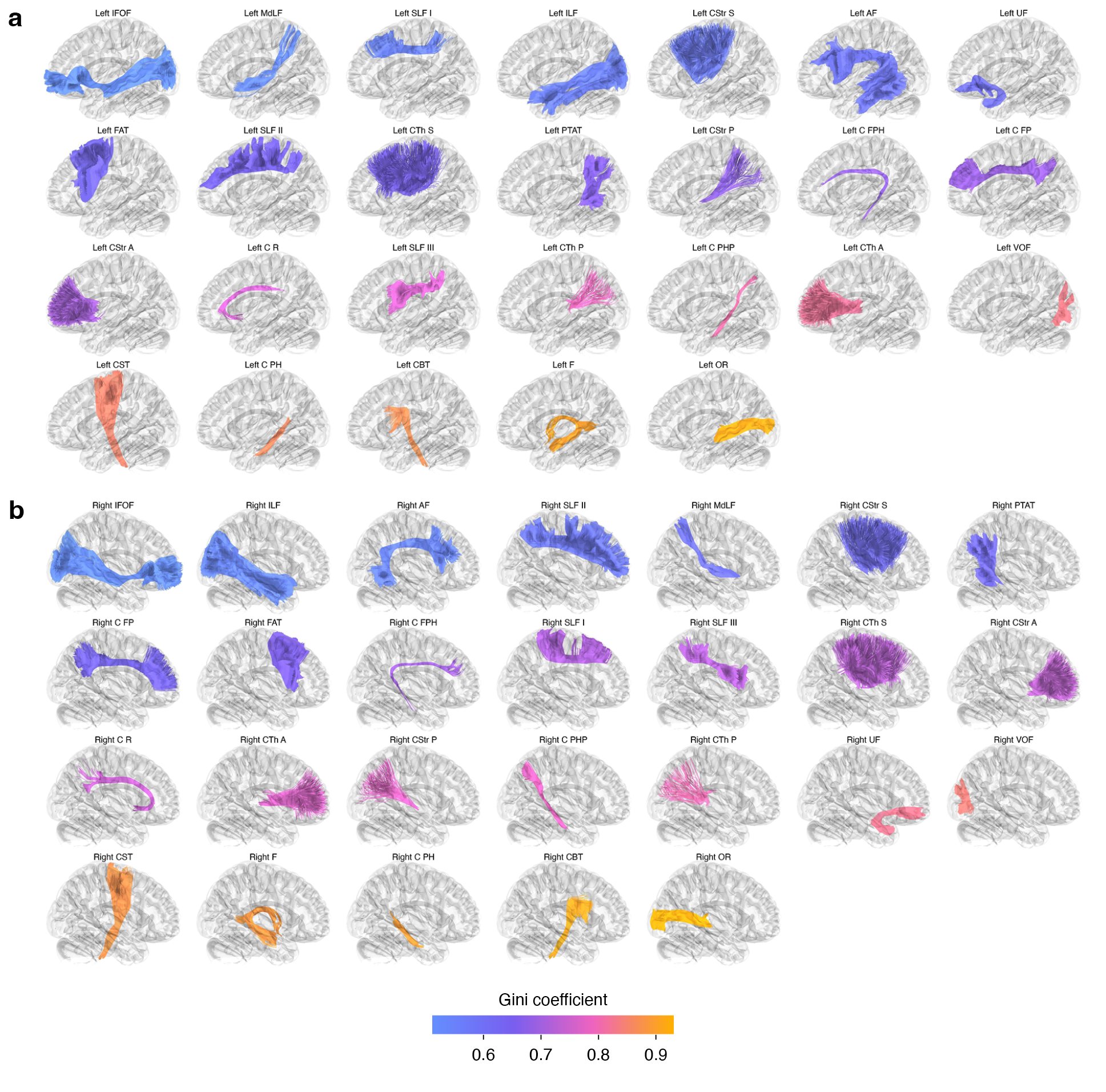

### **Supplementary Figure 2**. Gini coefficient of diversity in WM tracts.

Individual anatomical WM tracts are shown colored by their Gini coefficient of cognitive diversity. **a)** left hemisphere, **b)** right hemisphere. Table 1 of the manuscript contains tract abbreviations and full names for all 52 tracts.

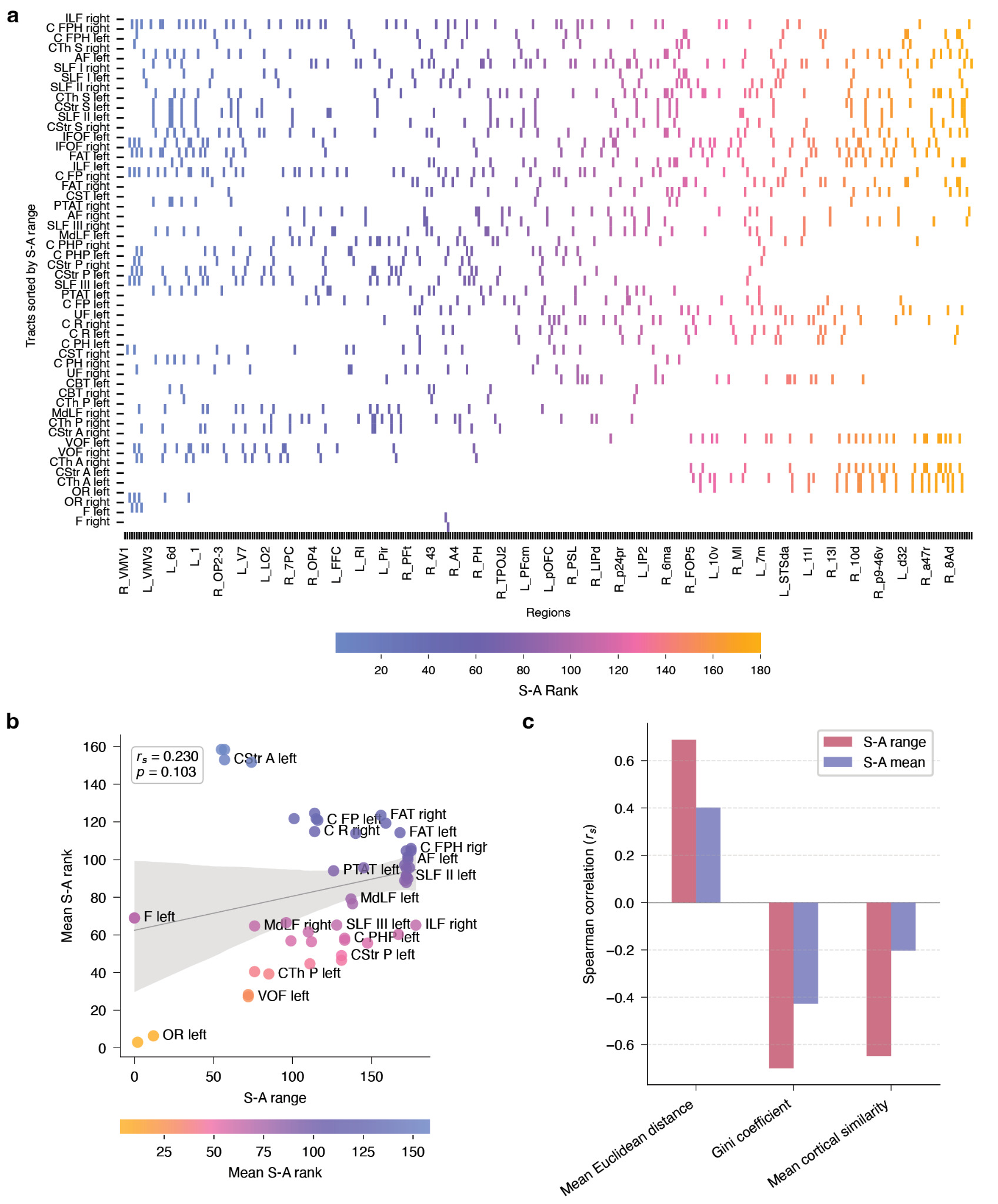

### **Supplementary Figure 3**. Measures of tract positioning in the cortical hierarchy.

**a)** Heatmap of the sensorimotor-to-association (S-A) hierarchy ranks of cortical regions connected to each white matter tract. Tracts are ordered by descending S-A range, revealing that long-range tracts show varying connection patterns across the cortical hierarchy, with some connecting widely across the hierarchy, and others covering more narrow ranges. **b)** Scatterplot comparing two different measures of a tract’s position in the cortical hierarchy. The S-A range quantifies how widely a tract spans the hierarchy (maximum minus minimum S-A rank across connected regions), whereas mean S-A reflects the average S-A rank across connected regions. Tract S-A range and mean S-A were not significantly correlated (*r_s_*=0.23, *p*_perm_=0.1), indicating that each measure captures different information about a tract’s hierarchical position. **c)** Barplot summarizing the Spearman correlations between tract S-A range and other tract features described in the paper, including mean Euclidean distance between connected regions, the Gini coefficient of cognitive diversity, and mean cortical similarity reflecting the biological homogeneity of connected regions. For comparison, the same correlations were computed using each tract’s mean S-A rank. Across features, S-A range showed consistently stronger correlations, indicating that a tract’s hierarchical span explains more variance and represents an informative measure of a tract’s positioning within cortical organization.
